# Computational evidence of a new allosteric communication pathway between active sites and putative regulatory sites in the alanine racemase of *Mycobacterium tuberculosis*

**DOI:** 10.1101/346130

**Authors:** Jayanthy Jyothikumar, Sushil Chandani, Tangirala Ramakrishna

## Abstract

Alanine racemase, a popular drug target from *Mycobacterium tuberculosis*, catalyzes the biosynthesis of D-alanine, an essential component in bacterial cell walls. With the help of elastic network models of alanine racemase from *Mycobacterium tuberculosis*, we show that the mycobacterial enzyme fluctuates between two undiscovered states—a closed and an open state. A previous experimental screen identified several drug-like lead compounds against the mycobacterial alanine racemase, whose inhibitory mechanisms are not known. Docking simulations of the inhibitor leads onto the mycobacterial enzyme conformations obtained from the dynamics of the enzyme provide first clues to a putative regulatory role for two new pockets targeted by the leads. Further, our results implicate the movements of a short helix, behind the communication between the new pockets and the active site, indicating allosteric mechanisms for the inhibition. Based on our findings, we theorize that catalysis is feasible only in the open state. The putative regulatory pockets and the enzyme fluctuations are conserved across several alanine racemase homologs from diverse bacterial species, mostly pathogenic, pointing to a common regulatory mechanism important in drug discovery.

**Author summary:** In spite of the discovery of many inhibitors against the TB-causing pathogen *Mycobacterium tuberculosis*, only a very few have reached the market as effective TB drugs. Most of the marketed TB drugs induce toxic side effects in patients, as they non-specifically target human cells in addition to pathogens. One such TB drug, D-cycloserine, targets pyridoxal phosphate moiety non-specifically regardless of whether it is present in the pathogen or the human host enzymes. D-cycloserine was developed to inactivate alanine racemase in TB causing pathogen. Alanine racemase is a bacterial enzyme essential in cell wall synthesis. Serious side effects caused by TB drugs like D-cycloserine, lead to patients’ non-compliance with treatment regimen, often causing fatal outcomes. Current drug discovery efforts focus on finding specific, non-toxic TB drugs. Through computational studies, we have identified new pockets on the mycobacterial alanine racemase and show that they can bind drug-like compounds. The location of these pockets away from the pyridoxal phosphate-containing active site, make them attractive target sites for novel, specific TB drugs. We demonstrate the presence of these pockets in alanine racemases from several pathogens and expect our findings to accelerate the discovery of non-toxic drugs against TB and other bacterial infections.

## Introduction

Tuberculosis is one of the top 10 causes of mortality globally and according to latest available estimates, 10.4 million people developed this disease in 2016, of which 4.9 million people were infected with multidrug-resistant TB strains (MDR-TB) [1]. The prevalence of multidrug-resistant TB (MDR-TB) and extensively drug-resistant tuberculosis (XDR-TB) necessitates the inclusion of novel anti-tubercular therapies and strategies in the treatment of TB. Treatment regimen comprising simultaneous use of multiple drugs is the current strategy in practice [2]. Despite the implementation of this strategy, TB mortality rates have not abated. Therefore, efforts to eradicate the TB pandemic have been stepped up globally through research oriented towards finding new drugs against the tubercle bacilli [3].

Alanine racemase (EC 5.1.1.1; Alr), an essential bacterial enzyme [4] is a popular drug target due to the absence of human homologs. The enzyme catalyzes the inter-conversion of L- and D-alanine and requires pyridoxal 5’-phosphate (PLP) as a cofactor. PLP is covalently attached to the enzyme through an internal Schiff’s base linkage [5]. In the L to D direction, the enzyme catalyzes the formation of D-alanine, an essential component of D-alanyl-D-alanine found in the peptidoglycan layer in bacterial cell walls [5]. In some bacteria including *Escherichia coli* [6], *Salmonella typhimurium* [7] and *Pseudomonas aeruginosa* [8], there are two Alr isozymes (Alr1 and Alr2 (aka DadX)), responsible for the anabolic and catabolic functions respectively.

The catalytically active form of Alr is a dimer [9], due to the participation of residues from both the monomers towards the formation of a functional active site. A narrow passage from the exterior forms an entryway to the substrate binding cavity in the active site and is lined by conserved residues, some of which have been demonstrated to orient the substrate molecules during their entry into the active site [10, 11]. In Alr*_Mtb_*, the substrate binding cavity is a small, conical space gated by two tyrosine residues (inner gates), which restrict the entry of substances into the active site [12]. Carboxylates such as acetate, propionate and substrate analogs such as alanine phosphonate co-crystallize in the substrate binding cavities of alanine racemases [13–15] and are suggested to regulate catalysis by competitive inhibition, though the exact control mechanisms are not known [16].

Including the structure of Alr*_Mtb_* [12], there are around a dozen and a half unique alanine racemase structures in protein databases [13, 17–23]. Though there has been considerable interest in elucidating the detailed catalytic mechanism of D- to L-alanine racemization in several organisms [5, 10, 24, 25], the regulatory aspects of catalysis suffer from lack of research. In spite of the discovery of a plethora of inhibitors against pathogenic Alr [26–28], only one of them has reached the market as a TB drug. This drug (D-cycloserine) is a structural analog of D-alanine and binds to all PLP-containing enzymes non-specifically, including those in the host, inducing toxic side-effects [29]. Current drug discovery efforts focus on finding safer, selective, non-substrate inhibitors. Several inhibitors of Alr are non-substrate leads, whose target sites on the enzyme are not known. Of these, five were shown to be non-toxic to mammalian cells in a high-throughput screen for anti-tubercular small molecule inhibitors [28]. Until now, there have been no studies concerning the binding sites of these five drug-like leads (Fig 1) on the enzyme. Considering the numerous hurdles in culturing *M. tb* and the urgency in developing novel drugs to contain the superbug strains, we sought to determine the target sites of these leads through computational studies.

**Fig 1.**
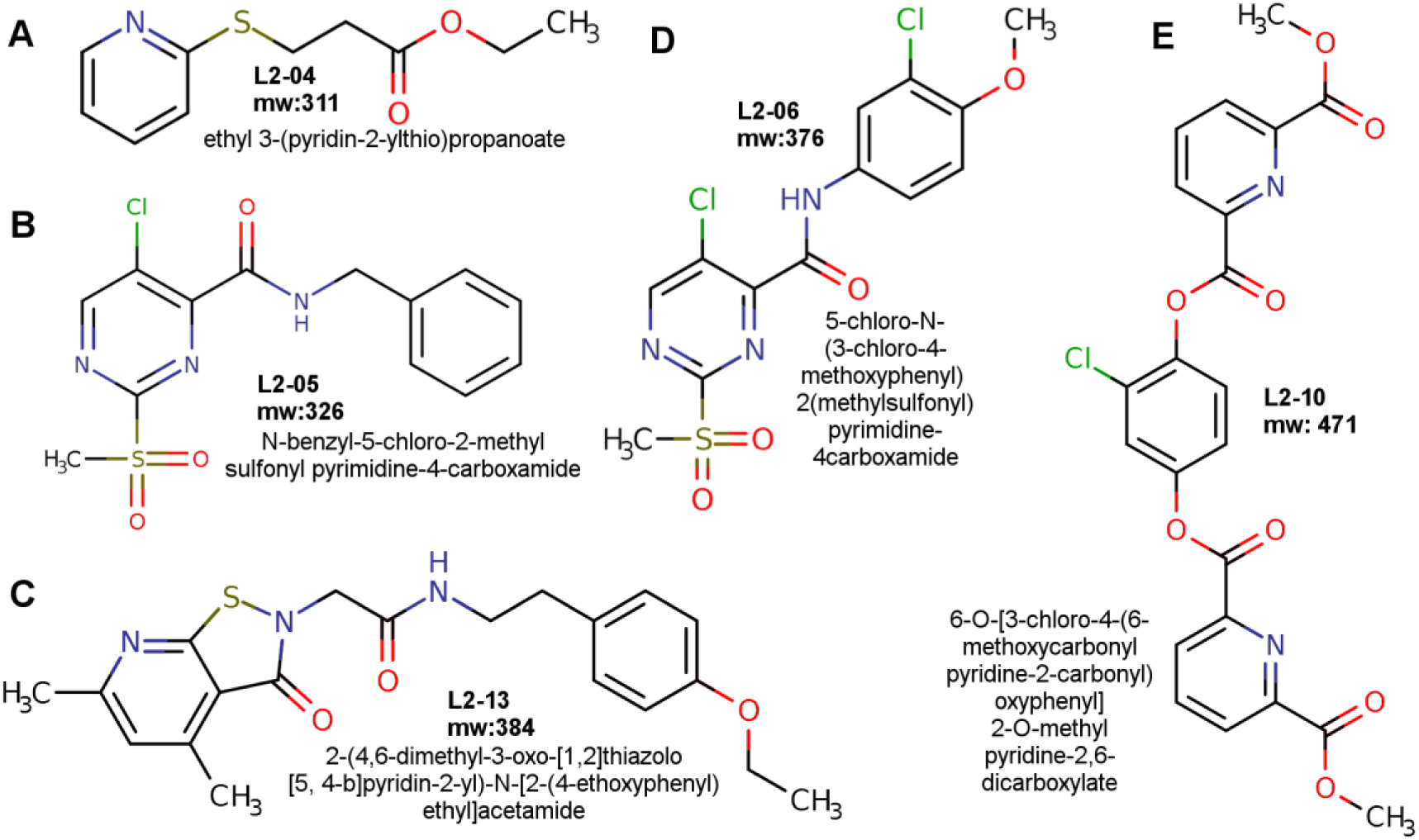
Inhibitors shortlisted from Anthony et al., 2011 [28]. IUPAC names and molecular weights are shown for each lead inhibitor. All of the listed inhibitors obey Lipinski’s ‘rule of 5’ characteristic of drug-like compounds and were shown to be non-cytotoxic to mammalian cells [28].

In recent years, normal mode analysis (NMA) has been widely used in probing large-scale, collective motions of proteins and has been increasingly utilized to characterize the dynamic aspects of enzymes [30–32]. Particularly, elastic network model (ENM) based NMA has been useful in studying intrinsic dynamics of slow protein motions over longer timescales [33, 34]. Computationally, the generation of elastic network models of diverse protein conformations is less expensive compared to molecular dynamics (MD) simulations [35]. In enzymes, ENM-NMA-predicted global motions represent biologically relevant functional motions and have been shown to include local fluctuations such as loop movements essential in catalysis [36]. We searched ENM-based Alr*_Mtb_* conformations for target sites of lead inhibitors through multiple, robust search algorithms by a blind docking strategy (BD). BD remains a common choice in the discovery of novel, allosteric binding sites [37, 38]. In conjunction with pocket search tools, BD is capable of identifying new functional pockets on the target protein [39]. This strategy helped us in the successful identification and validation of new pockets in Alr*_Mtb_*. Further to the above investigations, a comparative study of the intrinsic dynamics of Alr homologs with the help of a range of computational tools helped us gain new insights into the regulatory aspect of D-alanine synthesis.

## Results / Discussion

### All-atom normal mode analysis

#### The putative regulatory pockets are conserved across homologs

The crystal structure of Alr*_Mtb_* is a kidney-shaped dimer, with two active site cavities opening on the convex side (Figs 2A, 2C and 2E) and two pockets located on the concave side (Figs 2A and 2D). Residues found to be missing (Fig 2B) in the crystal structure were from both internal and terminal regions. The internal stretches of missing residues (176–180 of subunit A and 266– 280 of subunit B), pertained to the same region, i.e., the mouth region of the first active site cavity (Fig 2E).

**Fig 2.**
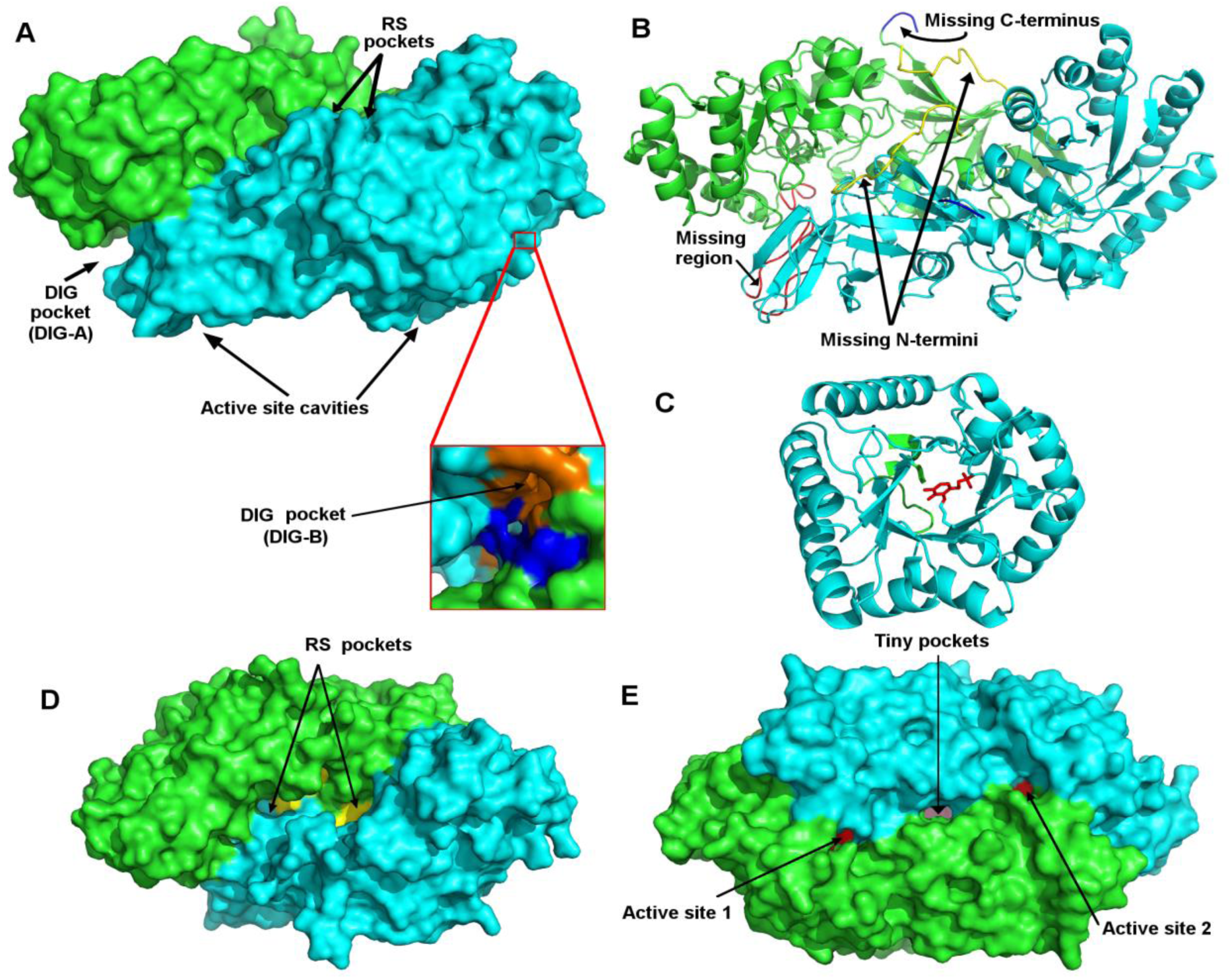
Structure of alanine racemase from *Mycobacterium* tuberculosis. A. Molecular surface representation of the structure of alanine racemase (monomers A and B shown in green and cyan colours respectively). Magnified region shows the putative dimer interface groove (DIG) pocket on the dimer interface. B. Unresolved regions in the crystal structure of Alr indicated by different colours in the cartoon representation of the enzyme (missing N-terminus—yellow; missing C-terminus—blue; missing internal stretches—red). C. TIM-barrel of active site 2 showing the cofactor PLP (red sticks) covalently attached to the catalytic residue Lys44 (green sticks). Note that the active site is composed of residues from both monomers (B monomer shown in cyan colour and residues from A monomer are coloured green) D. Surface representation of the enzyme showing the putative regulatory sites (yellow) E. Surface representation of the enzyme showing tiny pockets (pink) flanked on either side by the active site cavities (red). (Due to the revision in UniProt sequence information, the residue numbers given in this work should be decremented by 2 in order to compare with the numbering provided in LeMagueres et al., 2005 [12]. For example, the residues, 176–180 in our work refer to residues, 174–178 in LeMagueres et al., 2005 [12]).

Alignment of the protein sequences of Alr homologs (Figs 3, S1 and S2) revealed highly similar residues in the newly identified regions (described later): dimer interface groove region (Fig 3B), putative regulatory sites (Fig 3C) and a short helix (Fig 3D). On the other hand, the N-termini of the homologous Alr were of different lengths and were dissimilar in sequence composition (Fig 3A). Despite the presence of terminus in their sequences, 8 of the crystal structures of the homologs were devoid of either the N-terminus (varied between 3–15 residues) or the C-terminus (varied between 1–6 residues) or both. Of the remaining structures, eight were complete and showed disordered coils in their termini. Both PSI-PRED (secondary structure predictor based on position-specific-scoring-matrices of unique fold libraries) and Phyre2 (protein structure modeller based on a combination of *ab initio* and template-based strategies) generated highly disordered coils in the terminal regions (1–12 and 384–386) of *M. tb* racemase. Compared to the rest of the protein, the average B-factor values in the termini of most of the alanine racemases displayed a marginal increase. However, *Streptomyces lavendulae* (PDB ids: 1VFH, 1VFT, 1VFS), the closest structural homolog of Alr*_Mtb_* (49% sequence identity), displayed 2–3 times higher values in the N-terminal regions signifying a mobile terminus.

**Fig 3.**
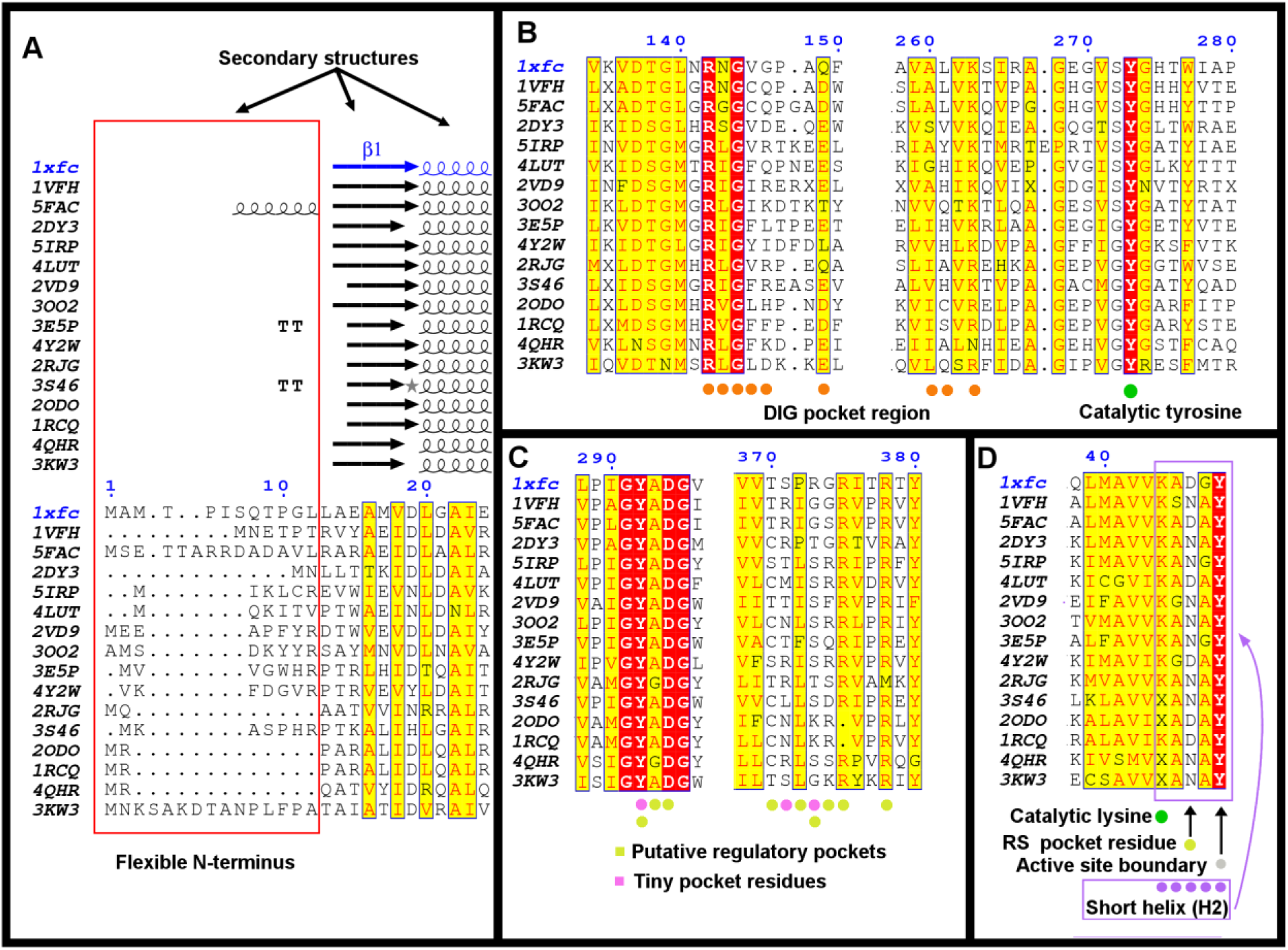
Structure-based multiple sequence alignment of alanine racemases. A. Aligned N-terminal region lacking definite secondary structures. B. Aligned DIG (dimer interface groove) pocket region. C. Aligned putative regulatory pocket and tiny pocket regions. D. Aligned region of the short helix (H2) residues implicated in the allosteric communication between the putative regulatory pockets and the active site residues. Shaded circles in different colours beneath the alignment show the residues in the corresponding pockets. (The alignment included sequences of *M. tuberculosis* (PDB code: 1XFC), *S. lavendulae* (PDB code: 1VFH), *S. coelicolor* (PDB code:5FAC), *C. glutamicum* (PDB code: 2DY3), *B. subtilis* (PDB code: 5IRP), *C. difficile* 630 (PDB code: 4LUT), *B. anthracis (Ames) (*PDB code: 2VD9*), S. aureus* (PDB code: 3OO2), *E. faecalis* (PDB code: 3E5P), *C. subterraneus subsp. tengcongensis* (PDB code: 4Y2W), *E. coli* (PDB code: 2RJG), *S. pneumoniae* (PDB code: 3S46), P*. fluorescens* (PDB code: 2ODO), *P. aeruginosa* (PDB code: 1RCQ), *A. baumannii* (PDB code: 4QHR), *B. henselae* (PDB code: 3KW3). X denotes cofactor ‘PLP-Lysine’ adduct, also called LLP).

Pockets were searched for in Alr*_Mtb_* as described in Methods after modeling the missing residues into its crystal structure. Pocket analysis of the Alr*_Mtb_* dimer model revealed the presence of two surface pockets beneath the terminal disordered coils. The pockets were adjacent to each other at the dimer interface, one corresponding to each subunit (Fig 2D). An examination of the crystal structures of the homologs revealed the existence of such pockets, either exposed to the exterior (PDB ids: 2RJG, 3E5P, 1RCQ, 2ODO) or sequestered beneath the terminal regions (PDB ids: 3S46, 2VD9, 4Y2W). Closed pockets sequestered beneath the termini were spacious enough to accommodate typical drug-like small molecules (combined Connolly’s molecular surface volumes of the two pockets were 358 Å^3^, 900.4 Å^3^, 834 Å ^3^ in 3S46, 2VD9, 4Y2W respectively). The inadvertent inclusion of extra diagonal spaces in the pocket volume calculations of Alr*_Mtb_* and the longer, lid-like N-termini (Fig 3A) resulted in a single larger pocket in *M. tuberculosis* (combined Connolly’s molecular surface volume of the two pockets in the closed state of Alr*_Mtb_* = 4355.9 Å^3^). In the current discussion, the mere presence of these pockets across species is of significance. Across homologs, two blocks of pocket residues (Fig 3C) were found to be conserved: region 292–295 in Alr*_Mtb_* (part of a conserved motif GY[AG]DG) and region 370–375 in Alr*_Mtb_* (375 is completely conserved, 372 is partially conserved). These regions were not exposed to the solvent in the crystal structures with sequestered pockets.

Upon examination of the normal modes of Alr*_Mtb_*, we observed that two low-frequency normal modes, LF_8_ and LF_10_ displayed the gradual closure of N-terminal regions over the surface pockets. Of the 30 conformations of LF_8_ (normal mode 8), 15 were non-identical and described the enzyme transition between an open state (conformations 8 and 9) and a closed state (conformations 23 and 24). The spectrum of conformations between the 9th and 23^rd^ showed the surface pockets at various stages of closure. Fluctuation plot of the normal mode 8 (Figs 4A and 4B) revealed that the close-open movements of the terminal regions are events, resulting from the closing and opening of the bulky N-terminal domains of the enzyme. In other words, the terminal regions always moved along with the other fluctuating regions in the mode 8 enzyme conformations. Such large-scale movements involving entire domains were observed in every examined homolog (shown as cumulative root mean square displacement (cRMS) plots in Figs 4C–4H). In the cRMS plots, fluctuation patterns of Alr were extremely similar in the bacterial genera belonging to the same phylum (Fig S3), viz., Actinobacteria (*M. tb* and *S. lavendulae*), Proteobacteria (*P.aeruginosa* and *P.fluorescens*) and Firmicutes (*Streptococcus pneumoniae* and *Enterococcus faecalis*).

**Fig 4.**
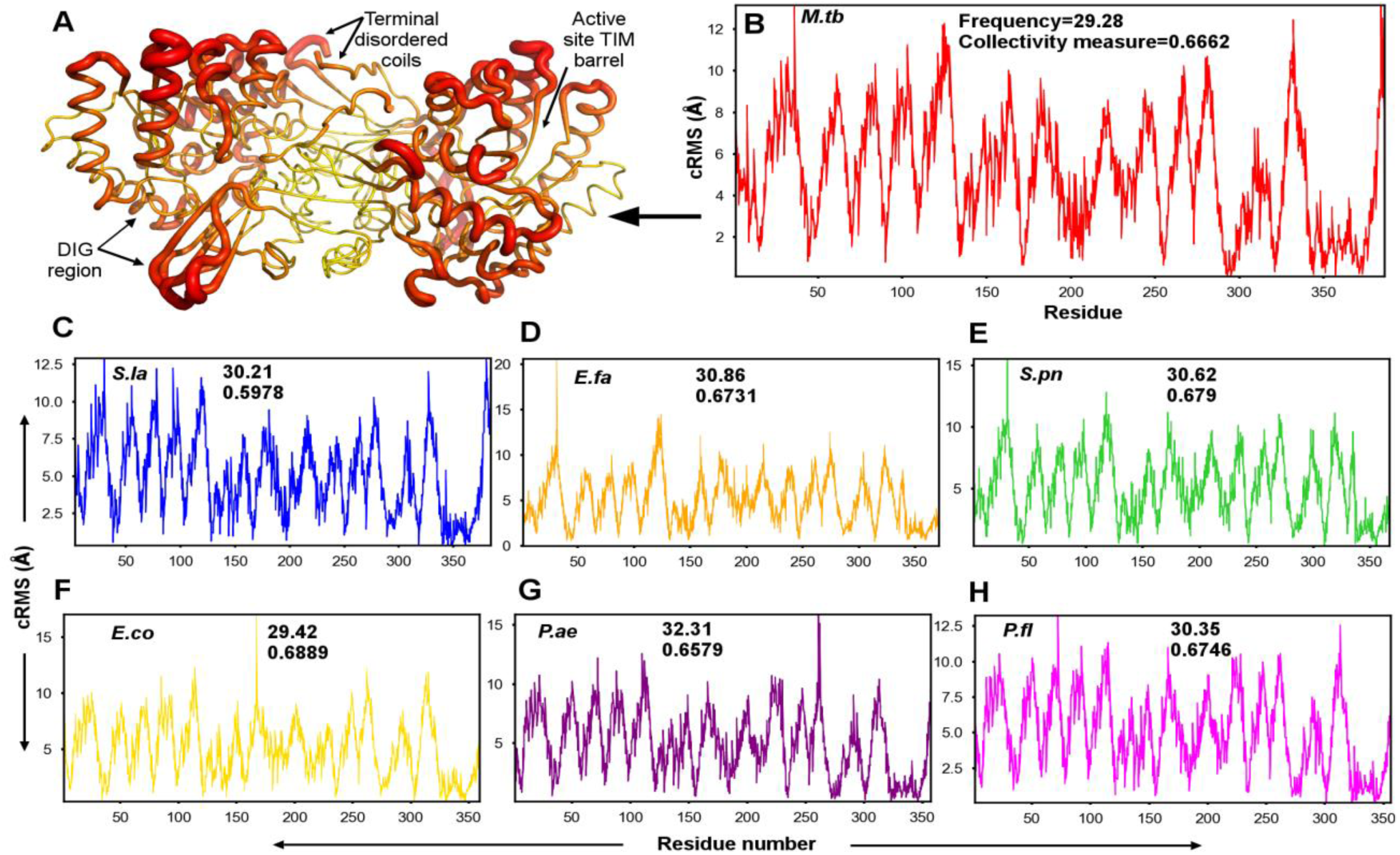
Fluctuations of bacterial alanine racemases. A. Putty cartoon view of fluctuations of Alr*_Mtb_* (Graph in panel B) mapped onto residues of the mycobacterial Alr structure colored from low to high values (0–13 Å as yellow to red). B–H. Plots of atomic displacements (derived from all-atom normal mode analyses calculated on NOMAD-Ref server) of bacterial alanine racemases, *M. tuberculosis* (PDB code: 1XFC), *S. lavendulae* (PDB code: 1VFH), *E. faecalis* (PDB code: 3E5P), *S. pneumoniae* (PDB code: 3S46), *E. coli* (PDB code: 2RJG), *P. aeruginosa* (PDB code: 1RCQ), *P. fluorescens* (PDB code: 2ODO), respectively along normal mode number 8 (LF_8_). Collectivity measures and frequencies of normal mode are indicated for each organism.

Overall, the fluctuation patterns, including the closure of the surface pockets (Movie S1) occurring as part of N-terminal domain movements were similar in all of the examined Alr homologs. In some of the homologs, the putative lid regions were shorter than those in *M*. *tb*. In the closed conformations of such homologs, the pockets were not completely covered.

Additionally, when the fluctuations of the first 10 normal modes were clustered (ensemble NMA based on C-α coordinates), higher amplitudes were observed in the termini of the homologs, *Streptomyces lavendulae* and *Caldanaerobacter subterraneus subsp. tengcongensis* (Fig 5). The conformational location of the terminus could be responsible for the observed oscillations, but that does not appear to be the case due to the following points:

1. Absence of definite secondary structures like α-helix or β-strand in the terminal regions of the structures of homologs.
2. Location of the disordered, flexible terminal regions over surface pockets in Alr homologs.
3. Presence of the pockets in one of the states in the crystal structures of homologs: ‘sequestered’ or ‘exposed’.
4. Closing and opening of the pockets during the intrinsic movements of the enzyme in normal mode simulations.
5. Docking of lead inhibitors to the surface pockets (discussed later).

**Fig 5.**
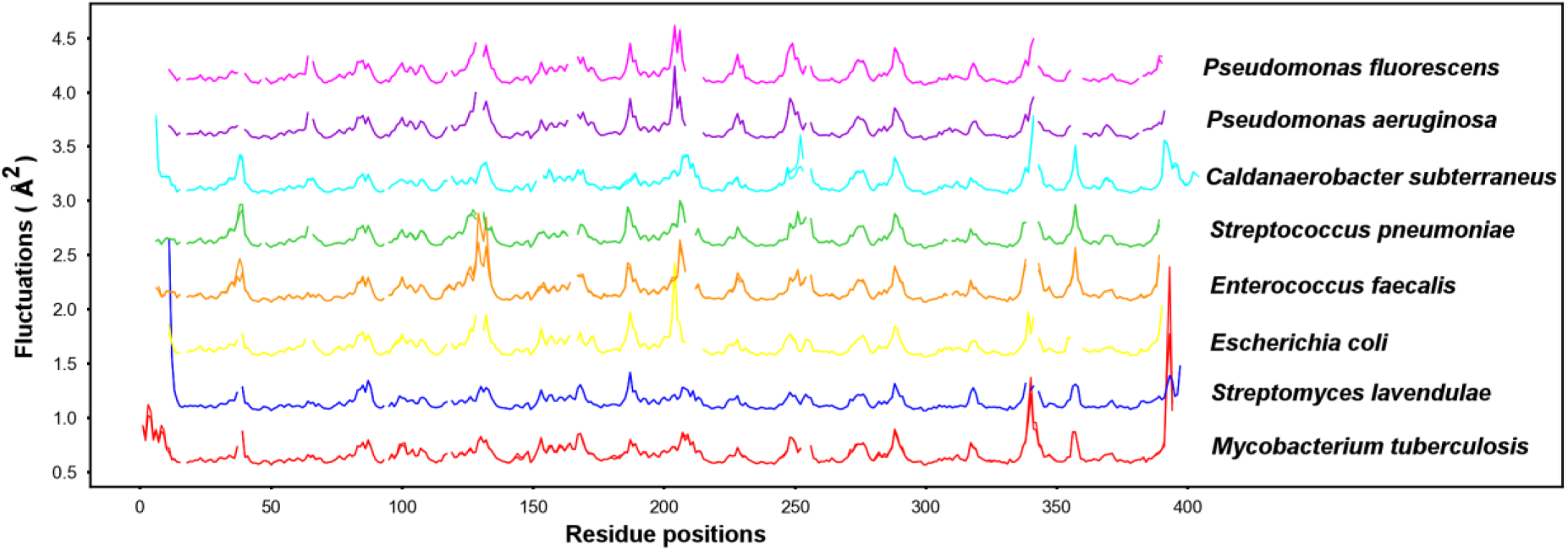
Cross species normal mode analysis of Alr. Ensemble normal mode analysis (eNMA) derived fluctuations (based on C-*α* coordinates) of the superimposed structures of the homologs. Superimposition was based on a multiple sequence alignment of protein sequences of Alr homologs on Bio3D modules, which was then manually edited for errors (Text S1). The fluctuation profiles of the Alr homologs are stacked at off-set values of 0.5 between each other on the y-axis for clarity. Gaps in the graphs correspond to the gaps in the aligned sequences.

Therefore, we conclude that the terminal fragments (especially the longer N-terminal 1–12 residue region in *M*. *tb*) anchored over the surface pockets help in sequestering small molecules, which are bound to the pockets. We propose that the two pockets in question are putative regulatory sites and refer to them as RS pockets (‘Regulatory Site’ pockets) throughout this study. As mentioned earlier, a few other regions in Alr*_Mtb_* fluctuated simultaneously with the close-open movements of the RS pockets:

1. Two new pockets (Fig 2A-magnified region) lying in the dimer interface junctions of the enzyme were found to elongate and contract with concomitant increase and decrease in their volumes. We name the pockets as DIG-A (Dimer Interface Groove pocket of monomer A) and DIG-B (Dimer Interface Groove pocket of monomer B).
2. Two tiny pockets lying adjacent to each other, in-between the entrances of the active site cavities (Fig 2E) exhibited close-open movements across the LF_8_ ensemble of conformations. The tiny pockets were high affinity binding sites of the native substrate (alanine) in both the crystal structure and the ensemble conformations. Charged residues, Arg373 and Glu367 of the tiny pockets were found to interact with the substrate. Across the ensemble conformations, the tiny pockets remained closed when the RS pockets were open. The tiny pockets and the RS pockets open on the opposite faces of the enzyme and share a few residues (292, 373) between each other. Other residues of the two pockets exhibit a unique arrangement in the sequence and are placed side by side in an alternating fashion (Fig 3C). Consequently, the adjacent residues in the structure belong to one of the two pockets and assume opposing states at any given instant during the dynamics. Therefore, the tiny and the RS pockets may be fulfilling opposing roles in regulation.

#### N-terminal domain movements lead to closure of regulatory pockets

Normal modes of frequencies less than 30 cm^−1^, cover most of the amplitudes in atomic displacements [40, 41]. A previous study has shown that a single normal mode is capable of carrying a lot of information on the conformational change of a given protein in terms of direction and pattern of atomic displacements [42]. In Alr*_Mtb_*, LF_8_ adequately described movements between two distinct states—one in which the putative RS pockets were open (Figs 6A, 6C) and the other in which they were closed (Figs 6B, 6D). Nearly 67% of amino acid residues were found to exhibit fluctuations (collectivity measure=0.6662). Among the all-atom normal mode ensemble of 30 conformations defining LF_8_, the closed and open conformations were both found twice each (conformations 8 and 9 represented open states while conformations 23 and 24 showed closed states). The pockets started closing from conformation 20 onwards and were fully closed in conformations 23 and 24. In the 23^rd^ and 24^th^ closed conformations, active site cavities were twisted from their original positions (Movie S1). A closer examination of Alr*_Mtb_* fluctuations showed that a twisted, hinge-like bending motion of the N-terminal domains of each monomer, transformed the open state into the closed state (Fig 6). The rigid body twist of the enzyme along the bulky N-terminal domains resulted in the closure of the putative N-terminal lid region over the RS pockets. The closed and open state fluctuations resembled the closing and opening of a door hinge with the two plates being the N-terminal domains and the centre of the hinge being the far end of the C-terminal region. Concordantly, higher deformation energies were seen in the pivot residues (96, 141,146, 261 and 263) of the dimer interface pocket region (DIG pocket region), part of which is the far C-terminal region (Fig 7). Apart from this hinge-like region, the putative terminal lid regions, the catalytic tyrosine and the short helix region showed higher deformation energy peaks signifying greater local flexibility (Fig 7).

**Fig 6.**
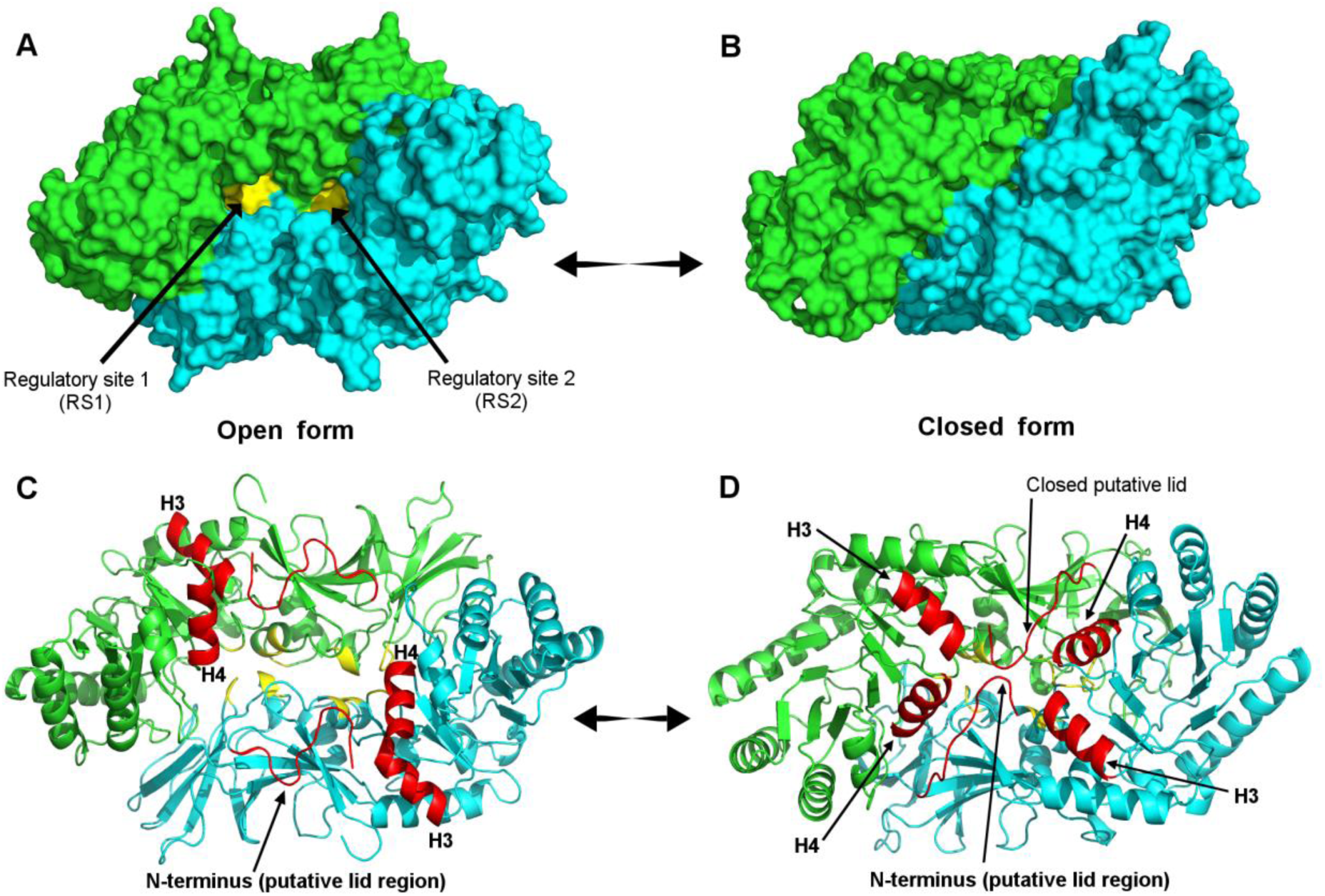
Depiction of transition of LF_8_ from an open to a closed conformation in Alr*_Mtb_*. (A) and (B). Representation of the molecular surfaces of the open and closed states of the mycobacterial alanine racemase. (C) and (D). Cartoon representations of the secondary structures of open and closed states of Alr*_Mtb_*. It can be noted that, the positions of helices (H3 and H4) have changed and the N-terminal putative lid-like regions are closed over the regulatory pockets in the closed state.

**Fig 7.**
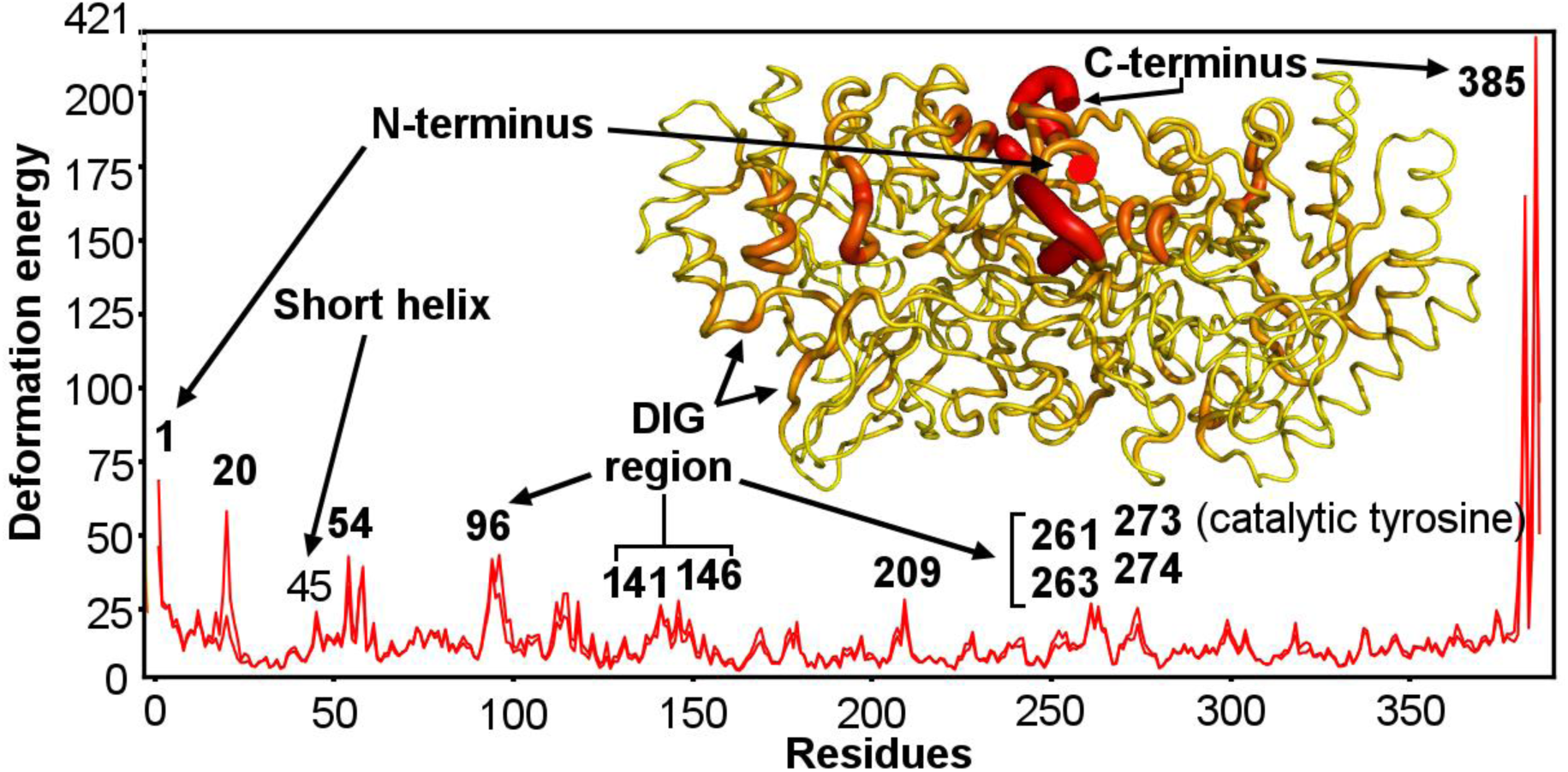
Visualization of mycobacterial Alr structure on the basis of local flexibility. A plot of residue-wise cumulative deformation energies derived from the first 10 low frequency normal modes of mycobacterial alanine racemase (energies calculated on C-α NMA of Alr*_Mtb_* on a standalone implementation of Bio-3D package). The putty view of deformation energies mapped onto mycobacterial Alr structure is coloured from low to high values (2–421 kcal / mol as yellow to red).

Table 1 portrays the loss of the relative orientation of the active site residues in space in the closed and open states as compared to that in the dimer model constructed from the crystal structure of Alr*_Mtb_* (See Methods). In the closed state of the active site 1, one of the inner gates, Tyr273, was 2.4 Å nearer to the other inner gate tyrosine, Tyr366, compared to their positions in the open state. As stated before, these two tyrosines guard the entrance of the substrate binding cavity. As discussed in LeMagueres et al. [12], these two residues define an opening of 2.7 Å in the crystal structure and are suggested by LeMagueres et al. [12] to move apart in order to permit entry and exit of small molecules in and out of the active site cavity. The distance between the two inner gates in the dimer model of crystal structure was 13.7 Å. This value is intermediate to the distance between the inner gates in the closed state (13.0 Å) and the distance between the same residues in the open state (15.4 Å). Thus, in the open state, the gates have moved further apart by 1.7 Å when compared to the distance between them in the model of the crystal structure. At the same time, the inner gates have come closer by 0.7 Å and are essentially closed in the closed state. Therefore, the active site entrance is closed in the closed state. The above results provide a proof for the hypothesis put forth by LeMagueres et al. that the inner gates must move apart prior to catalysis [12].

**Table 1.** Distances between C-α atoms of the substrate binding cavity residues^a^

| Residue range | Active site 1 |  |  |  | Active site 2 |  |  |  |
| --- | --- | --- | --- | --- | --- | --- | --- | --- |
|  | Distance between residues (Å) |  |  | Difference between open and closed states (Å) | Distance between residues (Å) |  |  | Difference between open and closed states (Å) |
|  | Open state (Conf. 8 and 9) | Closed state (Conf. 23 and 24) | Dimer model (Model of crystal structure with missing residues) (Conf. 16) |  | Open state (Conf. 8 and 9) | Closed state (Conf. 23 and 24) | Dimer model (Model of crystal structure with missing residues) (Conf. 16) |  |
| Tyr273 to Tyr 366 (Inner gates) | 15.4 | 13.0 | 13.7 | 2.4 | 14.8 | 12.6 | 13.3 | 2.2 |
| Tyr273 to Tyr48 | 19.3 | 18.4 | 18.1 | 0.9 | 18.7 | 18.1 | 17.6 | 0.6 |
| Tyr273 to Met321 | 12.2 | 11.4 | 11.3 | 0.8 | 11.5 | 10.9 | 10.7 | 0.6 |
| LYS44 to Tyr48 | 6.1 | 6.7 | 6.3 | 0.6 | 6.0 | 6.6 | 6.2 | 0.6 |
| Tyr273 to Lys44 | 18.7 | 18.7 | 17.8 | 0 | 18.1 | 18.2 | 17.2 | 0.1 |
| Cys320 to Tyr 366 | 10.0 | 9.5 | 9.5 | 0.5 | 10.1 | 9.6 | 9.6 | 0.5 |
| Met321 to Tyr 366 | 8.1 | 7.7 | 7.7 | 0.4 | 8.0 | 7.7 | 7.7 | 0.3 |
| Cys320 to Tyr48 | 12.6 | 12.9 | 12.4 | 0.3 | 12.7 | 13.1 | 12.5 | 0.4 |
| Met321 to Lys44 | 9.5 | 9.2 | 9.0 | 0.3 | 9.5 | 9.3 | 9.0 | 0.2 |
| Lys44 to Cys320 | 12.5 | 12.7 | 12.1 | 0.2 | 12.5 | 12.8 | 12.2 | 0.3 |
| Tyr273 to Cys320 | 8.6 | 8.3 | 8.0 | 0.3 | 8.0 | 7.9 | 7.5 | 0.1 |
| Met321 to Tyr48 | 9.1 | 9.3 | 8.9 | 0.2 | 9.2 | 9.4 | 8.9 | 0.2 |
| Lys44 to Tyr 366 | 12.0 | 11.9 | 11.6 | 0.1 | 11.9 | 11.9 | 11.6 | 0 |
| Tyr366 to Tyr48 | 7.4 | 7.5 | 7.2 | 0 | 7.4 | 7.5 | 7.2 | 0 |
| Arg142 to Met321 | 14.2 | 13.7 | 13.5 | 0.5 | 13.8 | 13.3 | 13.1 | 0.5 |
| Arg142 to Tyr366 | 21.0 | 19.4 | 19.6 | 1.6 | 20.7 | 19.2 | 19.4 | 1.5 |
| Arg142 to Tyr273 | 10.4 | 10.0 | 9.6 | 0.4 | 10.4 | 10.2 | 9.7 | 0.2 |
| Arg142 to Lys44 | 18.2 | 19.2 | 17.8 | 1.0 | 17.8 | 18.8 | 17.4 | 1.0 |
| Arg142 to Tyr48 | 21.6 | 22.1 | 21.1 | 0.5 | 21.2 | 21.7 | 20.7 | 0.5 |

<sup>a</sup>calculated on PyMOL molecular graphics tool

#### Ensemble NMA (eNMA) uncovers conservation of conformational dynamics across homologs

Comparative ensemble NMA (eNMA) results showed that flexibility profiles were largely similar across anabolic alanine racemases (PDB ids: 1XFC/*M*. *tb*; 1V FH/ *Streptomyces lavendulae*; 2RJG/*Escherichia coli*; 3E5P/ *Enterococcus faecalis*; 4Y2W/*Caldanaerobacter subterraneus subsp. tengcongensis*; 3S46/*Streptococcus pneumoniae*) as well as their catabolic counterparts (1RCO/*Pseudomonas aeruginosa*; 2ODO/*Pseudomonas fluorescens*) (Fig 5). Regions differing in amplitudes between the two types of racemases included alignment position 40 (higher amplitude in anabolic racemases) and alignment position 65 (higher amplitude in catabolic racemases). For the complete multiple sequence alignment and the alignment positions utilized in ensemble NMA of homologs, refer to Text S1. Across the homologs, the conserved residues constituting the invariant core of the enzyme were found clustered around the same amplitude (Fig S4-A). Moreover, the alignment positions displaying partial conservation of residues also fluctuated more or less to the same extent (Fig S4-B). Notably, conserved residue positions of the active site residues (alignment positions 51, 281 and 329 corresponding to Alr*_Mtb_* residues 48, 273 and 321 resp.), putative RS pockets (alignment positions 77, 302 and 383 corresponding to Alr*_Mtb_* residues, 74, 294 and 375 resp.), DIG region (alignment position 149 corresponding to Alr*_Mtb_* residue, 144), ‘Second cavity opposite to PLP’ (refer to subsection 2 of section 3 for details about this cavity) (alignment positions 140, 147, 179 and 330 corresponding to Alr*_Mtb_* residues 135, 142, 174 and 322) displayed nearly identical amplitudes across homologs.

In order to understand the nature of fluctuations between different regions of the enzyme, we calculated dynamic cross-correlation maps (DCCM) of positional fluctuations of amino acid residues (Fig 8). DCCM was based on the C-α coordinates of the first 10 normal modes of Alr*_Mtb_* as against the all-atom NMA results (discussed in the previous sections). Figs 8A–8C show that the correlations between residue pairs in Alr*_Mtb_* are in excellent agreement with the all-atom NMA results (Figs 4A and 4B). Additionally, DCCM provided insights into the directions of the movements between different regions. On the whole, the region between residues 134 and 199, moved in opposite directions with respect to the two flanking regions, 1–133 and 200–386. Both the completely correlated (Fig 8B) and completely anti-correlated (Fig 8C) regions revealed coupled networks of residues that play a role in the close-open transition of the enzyme. For example, strong correlations were seen in the following regions: putative lid region, catalytic lysine, RS pockets, tiny pockets and the delimiting boundary residues of the substrate binding cavity. Similarly, strong anti-correlations were seen between DIG residues (143, 144, 146 and 149), and the regions listed in Table 2. Majority of the residues of the ‘Second cavity opposite to PLP’ moved in opposite directions with respect to all the other regions of the enzyme. In agreement with the all-atom normal mode results, the inner gates of the substrate binding cavity moved with equal amplitudes in opposite directions. Thus, inner gates were moving away from each other during the transition of the closed state to the open state. Noticeably, the movement of the inner gates was occurring as a result of the movement and expansion of the entire active site. Thus the DCCM of Alr*_Mtb_* derived from C-α fluctuations forms an additional proof for the conclusion derived earlier from the all-atom NMA of Alr*_Mtb_* that the inner gates must move prior to catalysis.

**Fig 8.**
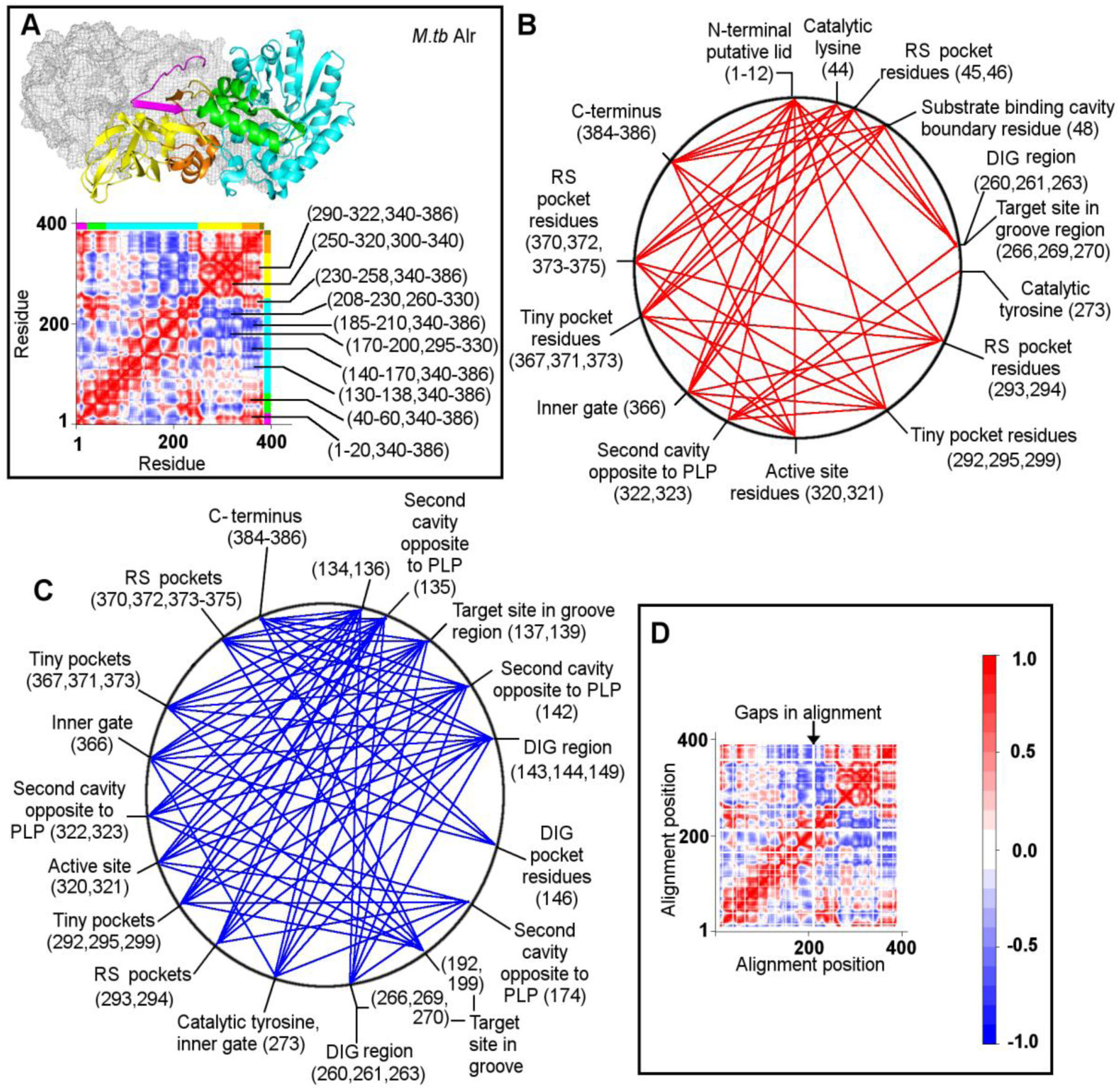
Correlation map revealing correlated and anti-correlated regions in Alr. A. Heatmap showing the dynamic cross correlation map of C-α atom fluctuations derived from an NMA of the first 10 low frequency modes of mycobacterial alanine racemase. Red regions denote the completely correlated residue pairs (same period, same phase) while blue regions denote the completely anti-correlated residue pairs (same period, opposite phase). White regions correspond to residue pairs whose fluctuations are not correlated. The top and right multi-colour bars on the heatmap correspond to various enzyme regions (coloured for easy visualization). B. Strongly correlated motions between mycobacterial Alr residues. Red lines portray the strong correlations between the regions mapped on the representative circle of a mycobacterial Alr monomer. C. Strongly anti-correlated motions between mycobacterial Alr residues. Blue lines portray the strong anti-correlations between the regions mapped on the representative circle of a mycobacterial Alr monomer. D. Dynamic cross correlation map of residual fluctuations derived from an ensemble-NMA of the first 10 low frequency modes of alanine racemases from 8 different bacterial species. Only those correlations present in all the 8 structures are shown in the plot.

**Table 2.** List of newly discovered target sites and their putative interactions with the inhibitors

| No. | Enzyme pocket | Ligand | Docking tools finding the enzyme-ligand complex | Enzyme residues involved in non-covalent interactions |
| --- | --- | --- | --- | --- |
| 1 | Putative RS pocket | L2-05, L2-06 | Vina, SwissDock | Ala45, Asp46, Glu74, Ala293, Gly374, Arg375, Arg378 |
|  |  | L2-10, L2-13 | Vina, SwissDock, PatchDock |  |
| 2 | DIG pocket region | L2-05, L2-06, L2-10 | Vina, PatchDock, SwissDock | Asp137, Gly139, Asn141, Gly146, Gln149, Arg192, Gln199, Lys263, Arg266, Glu269, Gly270 |
|  |  | L2-13 | Vina, SwissDock |  |
| 3 | Tiny pocket | Alanine | Vina | Tyr292, Gly295, Ser299, Glu367, Ser371, Arg373 |
| 4 | Second cavity opposite to PLP (cavity characterized in a previous study [12]) | L2-05, L2-06, L2-10, L2-13 | Vina, PatchDock | Trp90, Lys135, Arg142, His174, Asp322, Gln323 |

Both the correlation networks (Figs 8B and 8C) are each made of complete graphs with well-connected nodes. The highly connected nodes in both the networks are hub-like ‘hotspots’ which moved together either in the same or different directions. Therefore, perturbing any of the sites corresponding to these hotspot nodes, by targeting them with small molecules, would affect the regulatory control of catalysis, as all of these sites are involved in the transition of the enzyme from an open to a closed state.

In order to understand the pattern of correlations across homologs, we compared a residual fluctuation map showing only those correlations present in all the eight homologs (Fig 8D) with the individual DCCM of each homolog (Figs 8A, S5–S11). We found that the correlated and anti-correlated regions were nearly the same in all the maps, pointing to an evolutionary conservation of dynamics across alanine racemases. This conclusion is reiterated by the results of a comparative assessment of the normal modes of pairs of homologs using similarity measures of dynamics including RMSIP (Root Mean Square Inner Product) and Bhattacharyya coefficient (BC) (Table 3). RMSIP measures the similarity of atomic fluctuations derived from normal modes between proteins, whereas BC compares the covariance matrices obtained from the normal modes as elaborated in the Methods section. For example, a thermo-stable Alr from *Caldanaerobacter subterraneus subsp. tengcongensis*, which is a remote homolog of Alr*_Mtb_* (sequence identity=28.6%) shows 99.84% similarity in dynamics, as measured by the Bhattacharyya coefficient. Though the differences between the RMSIP scores were more pronounced than those of BC (Table 3), the latter is generally considered to be a better index for assessing the similarity of dynamics, as it incorporates eigenvalues. It is to be noted that RMSIP does not represent the energetic separation between the modes in the sets [43]. Sequence and structural similarity measures such as RMSD values scored lesser than dynamics similarity measures such as RMSIP and BC values (Table 3), proving that the conservation of dynamics far exceeds the sequence and structural conservation in alanine racemases.

**Table 3.**
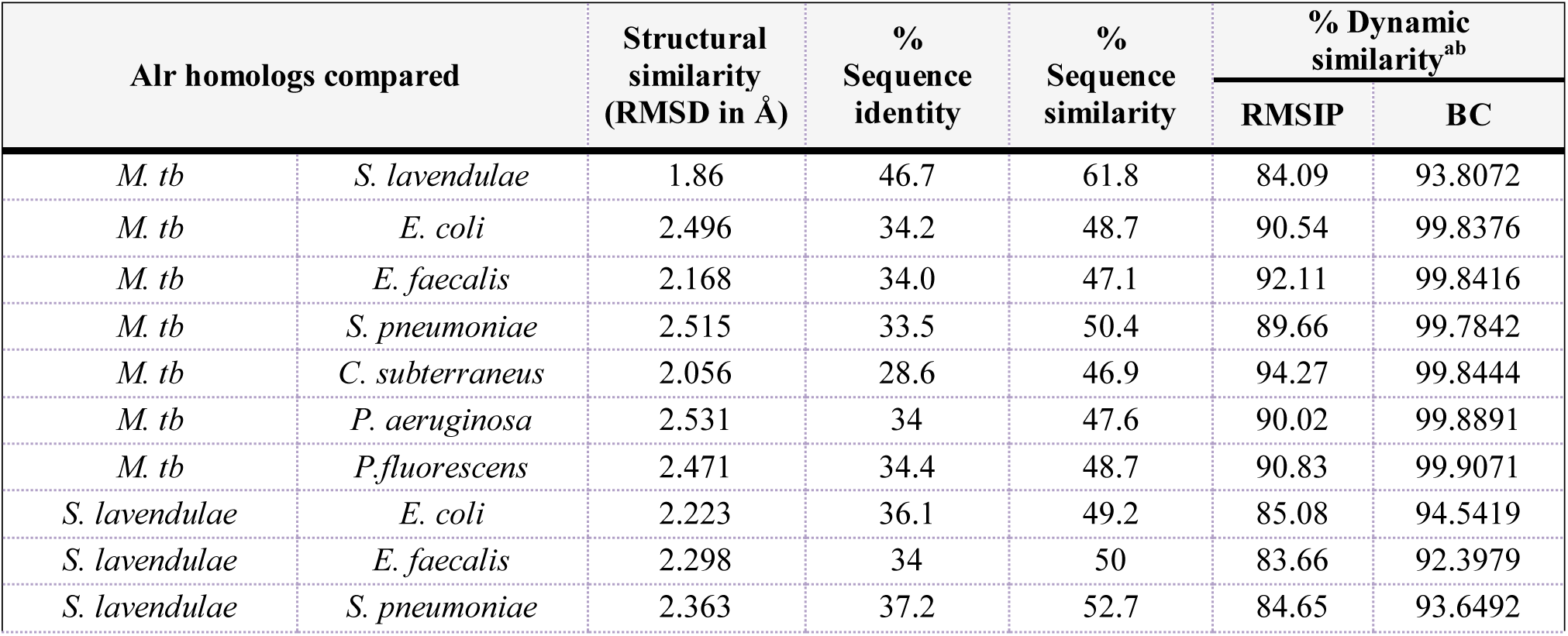
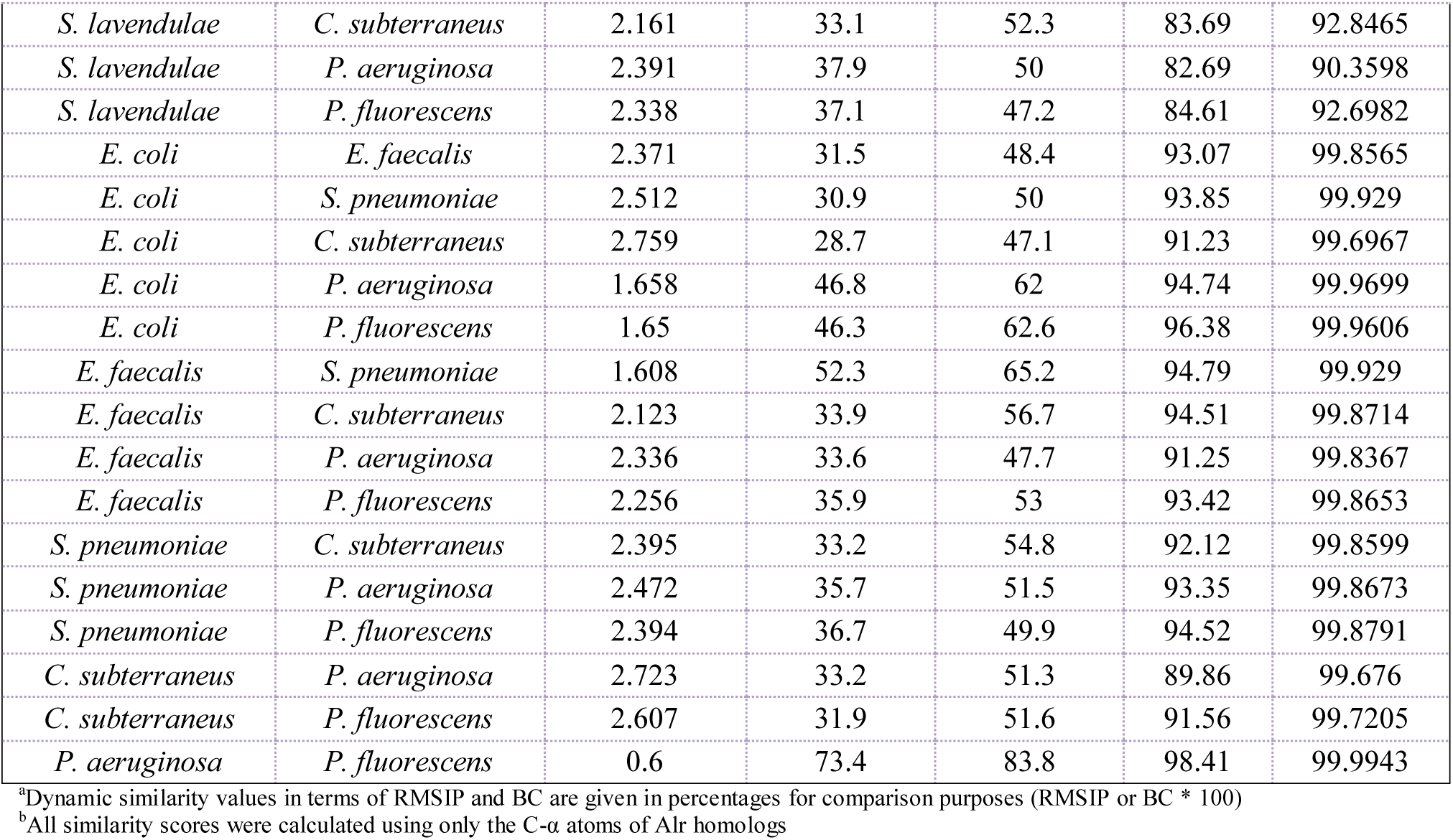
Measures of sequence, structural and dynamic similarity of alanine racemases across bacteria.

| Alr homologs compared |  | Structural similarity (RMSD in Å) | % Sequence identity | % Sequence similarity | % Dynamic similarity<sup>ab</sup> |  |
| --- | --- | --- | --- | --- | --- | --- |
|  |  |  |  |  | RMSIP | BC |
| *M. tb* | *S. lavendulae* | 1.86 | 46.7 | 61.8 | 84.09 | 93.8072 |
| *M. tb* | *E. coli* | 2.496 | 34.2 | 48.7 | 90.54 | 99.8376 |
| *M. tb* | *E. faecalis* | 2.168 | 34.0 | 47.1 | 92.11 | 99.8416 |
| *M. tb* | *S. pneumoniae* | 2.515 | 33.5 | 50.4 | 89.66 | 99.7842 |
| *M. tb* | *C. subterraneus* | 2.056 | 28.6 | 46.9 | 94.27 | 99.8444 |
| *M. tb* | *P. aeruginosa* | 2.531 | 34 | 47.6 | 90.02 | 99.8891 |
| *M. tb* | *P. fluorescens* | 2.471 | 34.4 | 48.7 | 90.83 | 99.9071 |
| *S. lavendulae* | *E. coli* | 2.223 | 36.1 | 49.2 | 85.08 | 94.5419 |
| *S. lavendulae* | *E. faecalis* | 2.298 | 34 | 50 | 83.66 | 92.3979 |
| *S. lavendulae* | *S. pneumoniae* | 2.363 | 37.2 | 52.7 | 84.65 | 93.6492 |
| *S. lavendulae* | *C. subterraneus* | 2.161 | 33.1 | 52.3 | 83.69 | 92.8465 |
| *S. lavendulae* | *P. aeruginosa* | 2.391 | 37.9 | 50 | 82.69 | 90.3598 |
| *S. lavendulae* | *P. fluorescens* | 2.338 | 37.1 | 47.2 | 84.61 | 92.6982 |
| *E. coli* | *E. faecalis* | 2.371 | 31.5 | 48.4 | 93.07 | 99.8565 |
| *E. coli* | *S. pneumoniae* | 2.512 | 30.9 | 50 | 93.85 | 99.929 |
| *E. coli* | *C. subterraneus* | 2.759 | 28.7 | 47.1 | 91.23 | 99.6967 |
| *E. coli* | *P. aeruginosa* | 1.658 | 46.8 | 62 | 94.74 | 99.9699 |
| *E. coli* | *P. fluorescens* | 1.65 | 46.3 | 62.6 | 96.38 | 99.9606 |
| *E. faecalis* | *S. pneumoniae* | 1.608 | 52.3 | 65.2 | 94.79 | 99.929 |
| *E. faecalis* | *C. subterraneus* | 2.123 | 33.9 | 56.7 | 94.51 | 99.8714 |
| *E. faecalis* | *P. aeruginosa* | 2.336 | 33.6 | 47.7 | 91.25 | 99.8367 |
| *E. faecalis* | *P. fluorescens* | 2.256 | 35.9 | 53 | 93.42 | 99.8653 |
| *S. pneumoniae* | *C. subterraneus* | 2.395 | 33.2 | 54.8 | 92.12 | 99.8599 |
| *S. pneumoniae* | *P. aeruginosa* | 2.472 | 35.7 | 51.5 | 93.35 | 99.8673 |
| *S. pneumoniae* | *P. fluorescens* | 2.394 | 36.7 | 49.9 | 94.52 | 99.8791 |
| *C. subterraneus* | *P. aeruginosa* | 2.723 | 33.2 | 51.3 | 89.86 | 99.676 |
| *C. subterraneus* | *P. fluorescens* | 2.607 | 31.9 | 51.6 | 91.56 | 99.7205 |
| *P. aeruginosa* | *P. fluorescens* | 0.6 | 73.4 | 83.8 | 98.41 | 99.9943 |

<sup>a</sup>Dynamic similarity values in terms of RMSIP and BC are given in percentages for comparison purposes (RMSIP or BC \* 100)
<sup>b</sup>All similarity scores were calculated using only the C-α atoms of Alr homologs

### Exploring inhibitor binding sites on Alr*_Mtb_*

Large-scale docking simulations (For details, see Methods) between the lead inhibitors (Fig 1) and the ensemble conformations (of the close-open transition) led to the following conclusions:

#### Substrate binding cavity is the primary target site of L2-04

L2-04 penetrated the substrate binding cavity in a majority of Vina and PatchDock complexes (122 Vina complexes; 103 PatchDock complexes (Table S1)). The aromatic ring side of L2-04 was often found in pi-stacking interactions between the inner gate residues, Tyr366 and Tyr273’ (residue labeled with a prime to indicate that it belongs to the opposite monomer) while its tail formed hydrogen bonds with the cofactor in the substrate binding cavity of Alr*_Mtb_*. Substrate binding cavity measures 5.5 × 5.0 × 2.5 Å^3^ and accommodates the substrate, L-alanine. Many guest substrates, substrate analogues and inhibitors such as acetate, propanoate, L-alanine phosphonate, lysine and D-cycloserine have been reported to occupy this cavity in homologs [13–15, 17, 44]. In the crystal structure of a thermo-stable Alr of a novel thermophile, *Caldanaerobacter subterraneus subsp. tengcongensis* [21], the substrate is found between the catalytic residues, Lys40 and Tyr268 (equivalent to Lys44 and Tyr273 in Alr*_Mtb_*) in the substrate binding cavity and forms hydrogen bonds (2.7 Å) with the catalytic tyrosine. We found that the substrate, alanine (Fig S12) and L2-04 docked similarly, into the active site of Alr*_Mtb_*, through hydrogen bonds with the hydroxyl ‘O’ atoms of the catalytic tyrosine, Tyr273’ (bond length:2.79 Å) and the cofactor PLP (bond length:2.91 Å). In a few complexes, L2-04 also formed salt bridges with the catalytic lysine (Lys44) of the enzyme. A superposition of the docked Alr—L2-04 complex with the crystal structures of Alr-ligand complexes from *Bacillus anthracis (Ames)*, *S. aureus*, *P. fluorescens* and *B. stearothermophilus* (Fig 9A) uncovered conserved enzyme residues interacting with the ligands across the complexes. Thus, the network of interactions of L2-04 with the active site residues, especially with the conserved catalytic residues, Lys44, Tyr273 and the cofactor PLP explains the effective abolition of racemization. Our results are in agreement with previous reports [28] which suggest that L2-04 binds Alr*_Mtb_* reversibly. We propose that L2-04 brings about reversible, competitive inhibition, where a competition set up between L2-04 and the substrate for the same space in the substrate binding cavity induces the establishment of a dynamic equilibrium between the two species. The relative concentrations of the substrate and L2-04, then decide the course of the catalytic process to proceed or terminate intermittently.

**Fig 9.**
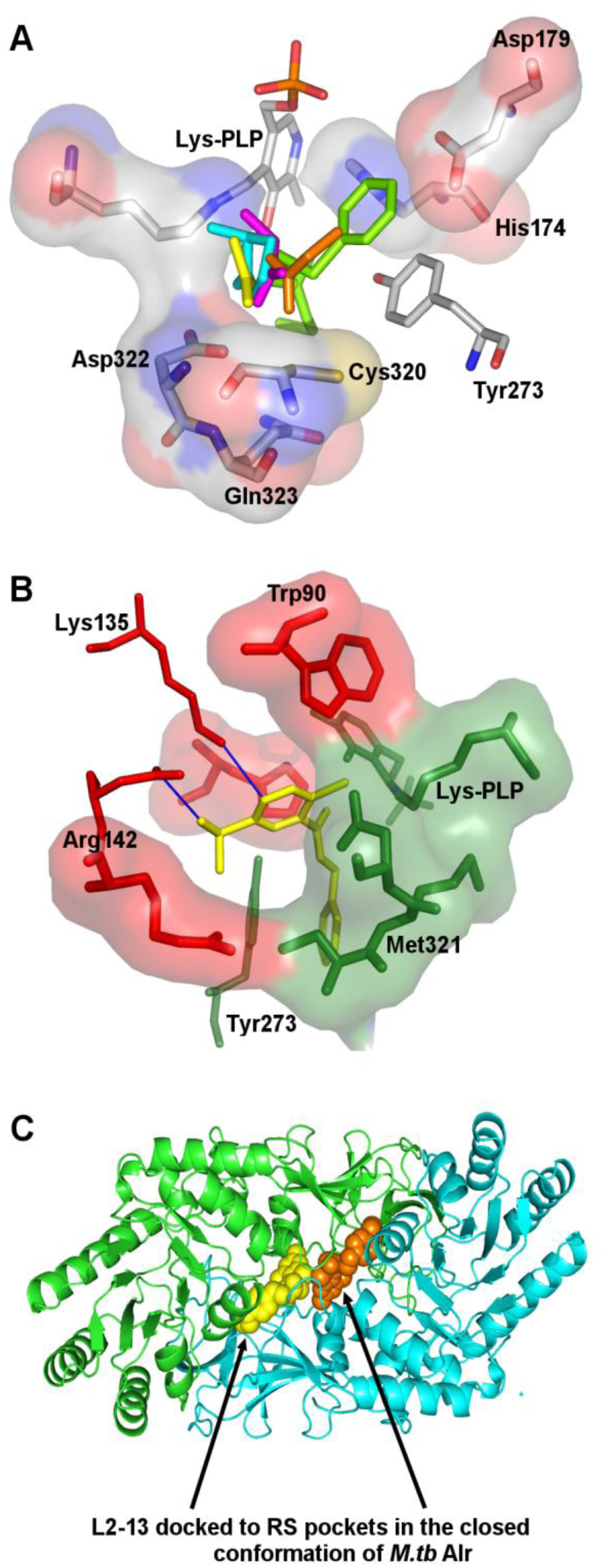
Docking of inhibitors to different pockets in Alr*_Mtb_*. A. Superposition of the substrate binding cavities of Alr with various bound ligands. Substrate binding cavity residues are shown as sticks and surfaces for reference. Ligands are represented as coloured sticks: docked L2-04 (green sticks); crystallized alanine (pink sticks) in Alr of *C. subterraneus subsp. tengcongensis*, PDB ID: 4Y2W; crystallized propanoate (orange sticks) in Alr of *E. faecalis*, PDB ID: 3E5P; crystallized 4-amino-isoxazolidin-3-one (cyan sticks) in Alr of *G. stearothermophilus*, PDB ID: 1XQL; crystallized acetate (yellow sticks) in Alr of *P. fluorescens*, PDB ID: 2ODO. B. L2-05 docked to active site in the closed state of Alr*_Mtb_* (Vina pose 1, Binding affinity: −8.5 kcal/mol; PatchDock poses 1 to 10). Note the entry of L2-05 (yellow sticks) into the ‘Second cavity opposite to PLP’ (cavity residues depicted as red sticks and surfaces) and the formation of hydrogen bonds (blue lines) with the residues Lys135, Arg142 in this cavity. Part of the inhibitor is still in substrate binding cavity (cavity residues depicted as green sticks and surfaces). C. L2-13 poses (yellow and orange spheres) docked to putative regulatory pockets in a cartoon representation of the closed form of Alr*_Mtb_*. (Orange spheres represent Vina pose 7 / PatchDock pose 7 docked to RS pocket corresponding to monomer B—binding affinity −8.2 kcal/mol; Yellow spheres represent Vina Pose 8 / PatchDock poses 3, 8 docked to RS pocket corresponding to monomer A—binding affinity −8.1 kcal/mol).

#### The target site extends into a cavity on the opposite side of PLP

Unexpectedly, a sizeable number of substrate poses bonded with residues of a second, larger (6.0 × 4.5 × 7.5 Å^3^) cavity adjacent to the active site on the opposite side of PLP. This cavity which has been previously characterized in *M*. *tb* by LeMagueres et al. [12] is accessible from the active site cavity. We refer to this cavity as ‘Second cavity opposite to PLP’ in this study. A very small number of high affinity poses of L2-05, L2-06, L2-10 and L2-13 were found to interact with the residues (Trp90, Lys135, Arg142 and His174) found in this cavity. Regardless of such interactions, none of the inhibitors were capable of occupying this cavity completely due to the presence of two or more ring systems in their bulky structures. Therefore, this cavity is less suitable for accommodating ligands of higher molecular weights, such as L2-05, L2-06, L2-10 and L2-13 (size range=370–470 daltons). Fig 9B shows L2-05 targeting a part of this cavity. Because the poses of all the 4 inhibitors are often found stretched between the active site and the second cavity, we believe that both the cavities are more suitable for ligands of lower molecular weight such as alanine (89 daltons).

#### Allosteric coupling to close-open movements are mediated by linker residues of a short helix

L2-05, L2-06, L2-10 and L2-13 (Fig 9C) targeted RS pockets multiple times through consensus pocket residues that included arginine, alanine and glycine. A consistent feature in all of the enzyme-inhibitor complexes was the interaction of the inhibitor with the pocket residues, Asp46 and Arg378, both of which were charged and placed adjacent to each other. Asp46 is situated on the same short helix (H2) of the active site TIM-barrel as the catalytic lysine (K44). While Lys44 is present on the inside of the active site TIM-barrel, Asp46 is present on the outside surface of the TIM barrel on the same short helix, but extending into the RS pockets.

Superposition of the open and closed states, both of whose RS pockets were bound with high affinity inhibitor poses (Fig 10A), clearly demonstrated the twisted active site cavity in case of the closed state. Docked poses of the bound inhibitors were observed to interact with charged RS pocket residues, Arg378 or Asp46 or both and such interactions appears to be driving the pull experienced by the short helix, H2 (Y48-G47-D46-A45-K44), seen in the normal mode motions. Such a movement of the short helix (Fig 10B) between the active site cavity and the RS pocket leads to the expansion and contraction of the active site cavities, as observed in the conformations of LF_8_. The catalytic residue, Lys44, linked by a covalent bond with the cofactor PLP on the inside of the TIM-barrels of the active site cavity (Fig 2C), would be dragged along with Lys44 towards the periphery of the active site cavity. As a result, the orientation of the catalytic residues would be lost. Tyr48, which walls the substrate binding cavity on one side through its side chain, forms the other end of the short helix and therefore would also be displaced, leading to the rearrangement of the substrate binding cavity (Fig 10B). Thus, the dynamic interactions between the inhibitor and the enzyme residues, viz., Arg378---Asp46 (R378---D46), is hypothesized to result in the short helix motion and aid in the communication between the regulatory and the active sites. This communication paradigm adequately explains the basis of signal (inhibitor binding to the regulatory site) transduction from the environment, all the way to the interior of the enzyme active site. The conservation of the charged arginine (Arg378) in the RS pockets across homologs (20 of the 21 homologs) point to a pivotal role for this residue in inhibitor interactions and regulation. In the short helix (YGDAK), the flanking residues, viz. the catalytic Lys44 and the active site residue, Tyr48 are completely conserved while the three middle residues showed equivalent substitutions (Glycine (G) by alanine (18/21), aspartic acid (D) by asparagine (14/21 times), Alanine (A) by either glycine (3/21) or serine (2/21)) (Fig 3D). A novel measure for amino acid flexibility in peptides devised by Huang et al. [45] ranks all the three middle residues of the short helix as highly flexible residues in the order, Glycine > Serine > Alanine, Aspartic acid and Asparagine. This result is in agreement with the need for higher conformational flexibility in the short helix residues in order to move between the RS pocket and the active site upon inhibitor binding. Supporting the above results, NMA studies show that deformation energies of the short helix residues are higher than the surrounding structure, indicating higher local flexibility (Fig 7). Generally, in TIM-barrel structures, there is a repetition of 8 alternating α helices and β strands. But, in case of alanine racemases, the arrangement of the active site TIM-barrel is as follows: α1-β1-α2-α3-β2-α4-β3-α5-β4-α6-β5-α7-β6-α8-β7-α9-α10-β8. It appears that the short helix H2 (α2), is an additional insertion (most likely by the splitting of the original second helix into α2 and α3) into the conventional TIM-barrel arrangement, the insertion event evolving probably later, in order to carry out allosteric regulation.

**Fig 10.**
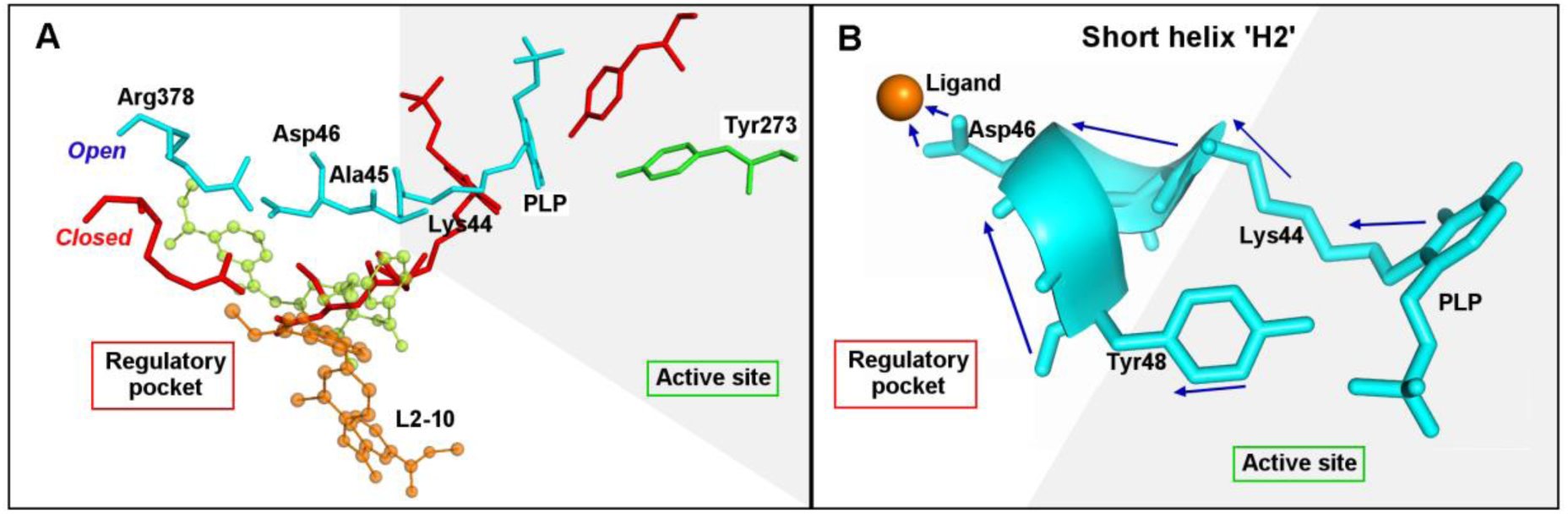
Distortion of active site geometry in mycobacterial Alr. A. Residue positions in the superimposed open and closed states. L2-10 docked to RS pockets in the closed state shown as orange ball and sticks (Vina pose 6 / PatchDock pose 10; Vina binding affinity: −9.1 kcal/mol); L2-10 docked to RS pockets in the open state shown as green ball and sticks (Vina pose 6 / PatchDock pose 5; Vina binding affinity: −8.3 kcal/mol). B. Depiction of the short helix (H2) residue displacements upon the binding of a hypothetical ligand. Tyr48 (Y48 delimits the substrate binding cavity on one side) and Lys44-PLP adduct (K44-PLP adduct important in catalysis) being pulled away from their original positions towards the periphery of active site cavity upon the binding of a representative inhibitor to Asp46, a regulatory pocket residue.

The structures of L2-05, L2-06, L2-10 and L2-13 are novel and not analogs of the substrate. Previous mass spectrometric analysis [28] have shown that three (L2-05, L2-06 and L2-13) of these inhibitors bind to the enzyme irreversibly. In these experiments, the binding mode of L2-10 could not be ascertained on account of an ambiguous peak profile. The irreversible binding displayed by the three (L2-05, L2-06 and L2-13) inhibitors in the mass spectrometry experiments when considered together with the docking results showing RS pocket-bound high affinity inhibitor poses, suggests that allosteric interactions of the inhibitors with the charged pocket residues (Asp46, Arg378) causes the irreversible inhibition of the enzyme.

#### Which of the enzyme states are catalytically active?

In the open state, the inner gates of the active site entrance were the farthest from each other and were completely open. In agreement, ensemble docking simulations showed that the substrate docked to the substrate binding cavity in the open state (Active site 1) and intermediate conformations but not in the closed state. In the open state, the substrate formed non-covalent interactions with the same set of conserved active site residues as seen in the crystal structure of the thermo-stable Alr (PDB ID: 4Y2W of *Caldanaerobacter subterraneus subsp. tengcongensis*) (Table 4). In contrast, the active site entrance was twisted and closed in the closed state, rendering the entryway (active site entrance) inaccessible. In such a shut active site cavity, catalysis is not feasible. Therefore, we reason that the open state is catalytically active.

**Table 4.** Hydrogen bonds between L-alanine and alanine racemase residues

| Structure | Type | Organism | Active site no. | Pose no. | Distance between L-alanine and PLP (Å) | Distance between L-alanine and Tyr273 or equivalent (Å) | Distance between L-alanine and Met321 or equivalent (Å) | Distance between L-alanine and Tyr292 or equivalent (Å) |
| --- | --- | --- | --- | --- | --- | --- | --- | --- |
| Crystal structure | Reference structure | *Caldanaerobacter subterraneus subsp. tengcongensis* | 1 | - | - | 2.67 | 2.56 | 2.8 |
|  |  |  | 2 | - | 2.91 | 2.96 | - | - |
| State 8 (open state) | Normal mode conformat | *Mycobacterium tuberculosis* | 1 | 10 | 2.86, 3.06 | - | 3.22 | - |
| State 10 | ions |  | 1 | 8 | 2.84,<br>3.09 | - | 3.19 | - |
| State 13 |  |  | 1 | 8 | 2.87,<br>3.03 | - | 3.12 | - |
| State 18 |  |  | 1 | 8 | 2.74 | 3.03 | 3.1 | - |
| State 19 |  |  | 2 | 8 | - | 3.00 | 2.89 | 2.96, 3.04 |
| State 20 |  |  | 2 | 9 | - | 2.99 | 2.94 | 2.93, 2.98 |
| State 21 |  |  | 2 | 9 | - | 2.97 | 2.99 | 2.93, 2.95 |

## Conclusion

Despite the increasing popularity of allosteric inhibitors as potential therapeutic agents [46], the structural basis of the mechanism behind allosteric enzyme inhibition remains virtually unknown. Alr*_Mtb_* has more than one active site as is characteristic of an allosteric enzyme. The presence of pockets at a location physically distant from the active site together with the fact that the pocket volumes are optimum to house molecules in the size range of natural regulators (activators or inhibitors), reinforces the theory of allostery. The closure of the putative regulatory pockets, occurring as part of native dynamics indicates that such an event is part of the catalytic process. Without the occupation of the pockets by natural modulators during catalysis *in vivo*, it would be needless for the pockets to be undergoing opening and closing movements. Moreover, docking of inhibitors to the putative regulatory pockets, as evidenced by the ensemble docking results, fortify the allostery proposal. Conservation of close-open enzyme movements across several Alr homologs and the presence of 2 distinct states (closed and open RS pockets) in the crystal structures of different organisms make a strong case for a generic allosteric mechanism of regulation in alanine racemases.

In conclusion, computational investigations on alanine racemases aided in identifying new, potentially druggable pockets on Alr*_Mtb_*, bringing it back into focus as a promising drug target. Many marketed TB drugs are orthosteric ligands and cross-react with eukaryotic PLP-dependent enzymes limiting their usefulness as effective drugs. A pharmacophore model based on the knowledge of scaffolds of the newly identified, potentially druggable pockets will open doors to the discovery of novel, specific, allosteric antimicrobials against *M. tb* with lesser side effects. The normal mode fluctuations between the two different enzyme states is a testable computational hypothesis for experimental studies aimed at deciphering the regulatory mechanism of alanine racemization. Correlation of the short helix (Y48-G47-D46-A45-K44) motion to inhibitor binding at the allosteric site detailed in this work offers a structural basis for allosteric site-to-active site communication in Alr*_Mtb_*. Inferences on the nature of inhibitor binding are in good agreement with preliminary mass spectrometric experimental findings by Anthony *et al*. [28]. Mutation studies of the conserved residues of the putative allosteric sites will help in determining the role and essentiality of these new sites in the regulation of catalysis. The presence of conserved residues in the putative regulatory pockets and the similarity in the fluctuation profiles of Alr homologs points to a mode of regulation common to several bacterial species. The disruption of this common regulatory mechanism by targeting the new pockets with a single inhibitor in diverse bacterial pathogens is a prospective tip for the discovery of broad-spectrum antibiotics in future drug discovery efforts.

## Methods

### Preparation of *M. tb* alanine racemase structure

The PDB [47] structure of mycobacterial alanine racemase (PDB id: 1XFC) was retrieved and analyzed. Missing internal and terminal residues in each of the monomers (10 N-terminal, 3 C-terminal residues) were constructed on Phyre-2 server [48] and a standalone version of Modeller (v. 9.16) [49]. The revision of sequence information as per the latest release of UniProt-K B [50] (UniProt i d: P9W QA 9), necessitated the splicing of 2 extra residues, ‘methionine’ and ‘alanine’ before the N-terminus prior to modelling. Initially, Modeller optimized the model by the variable target function method (VTFM) with conjugate gradients (CG). Subsequently, a preparatory molecular dynamics (MD) procedure with simulated annealing (SA) (300 steps each of heating the model *in vacuo* from 150K to 1300K and 1000 steps each of cooling the structure from 1300K to 300K) and a final, short conjugate gradient optimization (43 steps) were carried out by Modeller. An elaborate energy minimization of the model was additionally carried out by utilizing GROMACS [51] with AMBER-99SB* [52] forcefield on the MDWeb server [53] (500 steps of energy minimization of H atoms followed by 500 steps of energy minimization of the structure restraining heavy atoms to their initial positions with a force constant of 500kj / mol*nm^2^). The energy minimized model, thus obtained, was assessed on PDBSUM / PROCHECK [54, 55] and ERRAT [56] by analyzing Ramachandran plots [57] and other stereo-chemical parameters. Secondary structures of the missing termini were predicted on the PSIPRED v 3.3 webserver [58].

### Comparative analysis of alanine racemase homologs

A PSI-BLAST v.2.7.0 [59, 60] search for Alr homologs against the PDB database yielded 16 distinct PDB structures in the last iteration (e-value threshold = 0.0001) from 16 different organisms. Out of the 16, 14 were anabolic alanine racemases (Alr) and 2 were catabolic alanine racemases (Alr2/DadX). All the 16 sequences were aligned on the ENDscript 2.0 web server [61] using the in-built Clustal Omega tool [62]. Upon examination of the PDB structures of these 16 sequences, eight of them (*M*. *tuberculosis* (PDB code: 1XFC), *S*. *lavendulae* (PDB code: 1VFH), *E. faecalis* (PDB code: 3E5P), *E*. *coli* (PDB code: 2RJG), *P*. *aeruginosa* (PDB code: 1RCQ), *P*. *fluorescens* (PDB code: 2ODO), *S*. *pneumoniae* (PDB code: 3S46), *C*. *subterraneus subsp. tengcongensis* (PDB ID: 4Y2W)) were found to be completely resolved and were selected for all further analyses. Root Mean Square Deviation (RMSD) between the aligned C-α coordinates of the atom pairs of the corresponding Alr homologs were calculated in order to compare the similarity of native structures between any two homologs.

### All-atom normal mode analysis on individual Alr structures

Elastic network, Tirion-style normal mode models of alanine racemases from *M. tb* (dimer model of PDB id: 1X FC) and 6 completely resolved Alr homologs viz., those of *Streptomyces lavendulae* (PDB id: 1VFH), *Escherichia coli* (PDB id: 2RJG, 2RJH), *Streptococcus pneumoniae* (PDB id: 3S46), *Enterococcus faecalis* (PDB id: 3E5P, 3E6E), *Pseudomonas aeruginosa* (PDB id: 1RCQ), *Pseudomonas fluorescens* (PDB id: 2ODO) were generated on NOMAD-Ref web server [63] to explore collective, functionally relevant movements in the enzymes. The first 10 non-trivial, low frequency normal modes were generated for all atoms of the enzyme from each microorganism. As there is no provision currently to include cofactors in NMA calculations, PLP was excluded in the NMA calculations and was remodeled later into the NMA ensemble conformations prior to docking.

NOMAD-Ref calculates the modes by considering a highly simplified, quadratic potential energy between atoms that are assumed to be linked by a spring of universal strength. The atom-pairs linked by such a spring of an arbitrary elastic constant of 100 kcal / mol / Å^2^, were considered to be located <10 Å away in all the examined PDB structures. Eigen frequencies of the normal modes were generated after weighting all the interactions by exp (−(d_ij_(0)/d_0_)^2^), where d_0_ is the distance-weight parameter, which effectively introduced a smoother “cut-off” value than the original Tirion model. Negative eigenvalues were set to zero in the final output [63]. We obtained all our elastic models by applying a distance weight parameter value of 6.8 Å for the elastic constant and utilized the in-built Arnoldi iterative algorithm in order to diagonalize a sparse hessian matrix of n X n dimensions (for example: n=17238 in *M*. *tb*). The amplitude of the protein movement was controlled by fixing the average RMSD at 5 Å in output trajectories.

### Comparative normal mode analysis of Alr homologs

In order to study the fluctuations in the conserved regions of the racemases, ensemble-NMA elastic models of the superimposed invariant core of the enzyme across the 8 completely resolved Alr and DadX structures discussed before were generated on the eNMA module of a standalone implementation of Bio3D v. 2.1-1 package [64]. In general, for all downstream analyses, the data was filtered and only the first 10 low-frequency modes of each homolog were included in the calculations. In ensemble-NMA calculations, multiple sequence and structural alignment methods are utilized to analyse homologs. From these alignments, equivalent atom positions across structure ensembles were selected and normal mode vectors determined by calculating the effective force constant Hessian matrix K as,

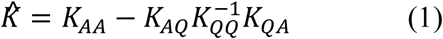

where K_AA_ represents the sub-matrix of K corresponding to the aligned C-α atoms, K_QQ_ for the gapped regions, and K_AQ_ and K_QA_ are the sub-matrices relating the aligned and gapped sites [65]. The normal modes of the individual structure in the ensemble can then be obtained by solving the eigenvalue problem,

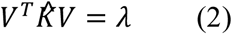

where V is the matrix of eigenvectors and λ, the associated eigenvalues.

In order to analyze the flexibility profile of the mycobacterial racemase, cross-correlations of residual fluctuations and deformation energy profiles were generated on the filtered normal mode data. Across homologs, the alanine racemase motions along the selected normal modes were compared with the help of similarity measures, viz., RMSIP and Bhattacharyya coefficient.

### Flexibility measures to assess Alr*_Mtb_*

#### Dynamic cross-correlation maps (DCCM)

The correlated motions undergone by C-α atoms of Alr*_Mtb_* protein during the first ten low-frequency normal modes were calculated using the Bio-3D DCCM module. The DCCM module generates a covariance matrix between residue pair fluctuations, i, j covering the entire length of the mycobacterial enzyme [66]. A cross-correlation coefficient is then calculated for each residue pair using the equation,

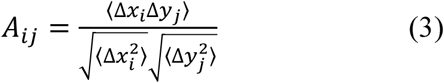

Here, Δ*x_i_* and Δ*y_j_* correspond to the displacement of i^th^ and j^th^ residues from their mean positions.

#### Deformation energy profiles

Deformation energies were calculated from raw Eigen energies and vectors of the first 10 normal modes of Alr*_Mtb_* (NMA based on C-α atoms) with the help of Bio-3D modules. Deformation energy is a normalised measure of the energy contributed by individual atoms of the model towards deformations of the structure [67].

### Similarity measures for comparison of enzyme motion across homologs

#### Root mean square inner product (RMSIP)

For comparing sets of normal modes, the root mean squared inner product (RMSIP) of the first 10 low frequency modes are generally included [68]. The RMSIP quantifies the similarity of the directions of these low energy subspaces (the subset of low frequency normal modes) between any two given proteins. RMSIP measures the cumulative overlap between all pairs of the ‘l’ largest eigenvectors, and is defined as:

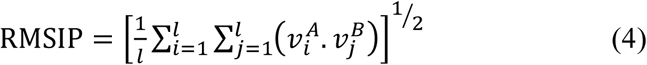

where 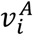 and 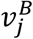 represent the i^th^ and j^th^ eigenvectors obtained from protein A and B, respectively, ‘l’ is the number of modes included (normally 10). The RMSIP measure varies between 0 (orthogonal) and 1 (identical directionality).

#### Bhattacharyya coefficient

The Bhattacharyya’s coefficient (BC) provides a means to compare two covariance matrices derived from NMA of two given proteins. For ensemble normal modes, the covariance matrix (C) can be calculated as the pseudo inverse of the mode eigenvectors:

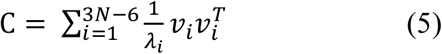

where *v_i_* represents the i^th^ eigenvector, *λ_i_* the corresponding eigenvalue, and N, the number of C-α atoms in the protein structure (3N−6 non-trivial modes). As formulated by Fuglebakk et al. [69], the Bhattacharyya coefficient can then be written as,

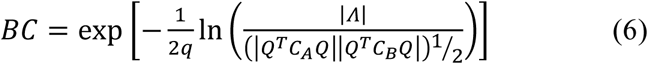

where Q is a matrix in which columns are eigenvectors of the averaged covariance matrix (*C_A_* + *C_B_*)12, Λ is a diagonal matrix containing the corresponding eigenvalues, and q the number of modes needed to capture 90% of the variance (cumulative eigenvalues) of Q. The BC varies between 0 and 1, and equals to 1 if the covariance matrices, *C_A_* and *C_B_* are identical.

### Docking simulations on the NMA ensemble conformations of Alr*_Mtb_*

Five potential drug-like inhibitors (Fig 1) from Anthony et al. (2011) [28] were short-listed and subjected to docking simulations against the LF_8_ ensemble of Alr*_Mtb_*. Prior to ensemble docking, the structures of the small molecules were prepared as elaborated below: 3D structures of the inhibitors were analyzed for all possible tautomers, stereoisomers, and major microspecies at pH 7 on the Chemicalize web server [70]. Lowest energy conformers were generated on a standalone implementation of Chemaxon MarvinView v.16.3.7.0 [71] by applying a Dreiding forcefield [72] at a diversity limit of 0.1 for a time limit of 1000s. L-alanine, the native substrate of the enzyme was included in the ligand set.

In an attempt to achieve higher accuracy, we employed AD Vina v.1.1.2 [73], which has an advanced scoring function and conducts a more exhaustive sampling of the possible binding modes. To minimize false positives, we applied two other docking algorithms with diverse search methodologies, viz., SwissDock [74] and PatchDock [75] for validation purposes. Although all the 3 docking algorithms employed robust methodologies for searching as well as scoring, we based our analyses primarily on AD Vina results. This strategy was adopted to nullify the bias exhibited by PatchDock and SwissDock towards active site cavities (PatchDock favoured active sites over other locations, while SwissDock clusters seldom entered active sites).

In addition to flexible inhibitors, we sought to introduce an implicit flexibility in the enzyme, by utilizing an ensemble of conformations, which sampled relevant functional movements. For this purpose, we selected 15 non-redundant conformations (9^th^ to 23^rd^ describing the ‘close-open’ transition) out of the 30 conformations defining the second low frequency mode 8 (LF_8_) of *M. tb* Alr, for all analysis. A total of 900 Alr-ligand complexes describing 10 best binding poses for each enzyme-ligand pair resulted from AD Vina runs. These results were validated by checking for identical target sites across 900 PatchDock complexes and approximately 3000 SwissDock clusters of binding poses with significant energies.

In general, the input structures were prepared according to the specifications of the docking software, retaining default values for all input parameters. In case of AD Vina, exhaustiveness factor was increased proportionally in view of the larger volumes of the search spaces probed, because of which, a minimum of around 100 runs were ensured for each enzyme-ligand combination. A standalone version of AutoDock Tools 1.5.6 [76] was used to prepare receptor and ligand structures in all AD Vina runs. AD Vina searches for ligand poses with the help of an iterated local search global optimizer algorithm and scores the results by an empirical X-score like function incorporating knowledge-based potentials. PatchDock finds docking transformations with good shape complementarity and ranks them based on geometric fit and atomic desolvation. SwissDock employs the EADock dihedral space sampling algorithm and incorporates CHARMM22 [77] forcefield energies by way of a multi objective scoring function using the FACTS solvation model [78]. Interactions of the docked ligands resulting from all of the simulation runs were visualized on PLIP v.1.3.2 [79] and an open-source version of PyMOL v.1.8 [80] (built and compiled from source code downloaded from SourceForge [81]).

### Pocket analysis

Normal mode conformations were investigated on the CASTp 3.0 [82] web server for the presence of pockets. Consensus enzyme residues targeted by inhibitors across pockets were gleaned by comparing the ligand-enzyme complexes with LigPlot+ v. 1.4.5 [83]. For all other input parameters, default values were retained. In general, custom perl scripts were used for all file processing purposes.

## Acknowledgments

We acknowledge the open source efforts in facilitating the distribution of vital molecular visualization tools like PyMOL as a freeware and would like to thank the open source software community and the developers in general.

## Author contributions

JJ designed and performed the computations and analysis. SC, TRK provided inputs and supervised, administered the entire project. JJ wrote the original draft of the manuscript while SC, TRK reviewed and revised the MS.

## Supporting Information

**Figure S1.** Multiple sequence alignment of Alr homologs from different bacterial species.

**Figure S2.** Sequence conservation profile of Alr*_Mtb_.*

**Figure S3.** Dendrogram based on the identity of protein sequences of alanine racemases.

**Figure S4.** A. Scatter plot of the normal-mode derived fluctuations of conserved residues of the superimposed homologs. This residue set corresponds to the invariant core of Alr*_Mtb_*. B. Scatter plot of the normal-mode derived fluctuations of similar, structurally equivalent residues of the superimposed homologs. All calculations were performed using C-α coordinates of the superimposed invariant core of alanine racemase homologs on the ensemble NMA (eNMA) Bio-3D module.

**Figure S5.** Dynamic cross-correlation map of residual fluctuations of alanine racemase of *Streptomyces lavendulae*.

**Figure S6.** Dynamic cross-correlation map of residual fluctuations of alanine racemase of *Enterococcus faecalis*.

**Figure S7.** Dynamic cross-correlation map of residual fluctuations of alanine racemase of *Streptococcus pneumoniae.*

**Figure S8.** Dynamic cross-correlation map of residual fluctuations of alanine racemase of *Caldanaerobacter subterraneus subsp. tengcongensis*.

**Figure S9.** Dynamic cross-correlation map of residual fluctuations of alanine racemase of *Pseudomonas aeruginosa*.

**Figure S10.** Dynamic cross-correlation map of residual fluctuations of alanine racemase of *Pseudomonas fluorescens*.

**Figure S11.** Dynamic cross-correlation map of residual fluctuations of alanine racemase of *Escherichia coli*.

**Figure S12.** LigPlot comparison of the docked substrate pose (L-alanine) of Alr*_Mtb_* with homologous Alr complexes co-crystallized with alanine in *Caldanaerobacter subterraneus subsp. tengcongensis*.

**Table S1.** Results of docking simulation runs of substrate and inhibitors on the ensemble conformations of alanine racemase from *Mycobacterium tuberculosis*.

**Table S2.** Properties of pockets of NMA ensemble conformations as calculated on the CASTp server.

**Text S1.** Multiple sequence alignment of Alr homologs utilized in ensemble NMA (shown with alignment positions).

**Movie S1.** Secondary structure representation of normal mode number 8 in Alr*_Mtb_*. N-terminal putative lid-like region is shown in red colour. Helices H3 (yellow) and H4 (violet) undergo displacements.

## References

1. World Health Organization [Internet]. World Health Organization. 2018 [cited 20 January 2018]. Available from: http://www.who.int/en/

2. Dheda K, Chang K, Guglielmetti L, Furin J, Schaaf H, Chesov D et al. Clinical management of adults and children with multidrug-resistant and extensively drug-resistant tuberculosis. Clinical Microbiology and Infection. 2017; 23(3):131–140.

3. Kumar V, Patel S, Jain R. New structural classes of antituberculosis agents. Medicinal research Reviews. 2017; 38(2):684–740.

4. Awasthy D, Bharath S, Subbulakshmi V, Sharma U. Alanine racemase mutants of *Mycobacterium tuberculosis* require D-alanine for growth and are defective for survival in macrophages and mice. Microbiology. 2011; 158 (2):319–327.

5. Watanabe A, Yoshimura T, Mikami B, Hayashi H, Kagamiyama H, Esaki N. Reaction mechanism of alanine racemase from *Bacillus stearothermophilus*. Journal of Biological Chemistry. 2002; 277 (21):19166–19172.

6. Wild J, Hennig J, Lobocka M, Walczak W, Klopotowski T. Identification of the dadX gene coding for the predominant isozyme of alanine racemase in *Escherichia coli* K12. MGG Molecular & General Genetics. 1985; 198(2):315–322.

7. Wasserman SA, Walsh CT, Botstein D. Two alanine racemase genes in *Salmonella typhimurium* that differ in structure and function. J Bacteriol. 1983; 153(3):1439–50.

8. Strych U, Huang H, Krause K, Benedik M. Characterization of the alanine racemases from *Pseudomonas aeruginosa* PAO1. Current Microbiology. 2000; 41(4):290–294.

9. Strych U, Benedik M. Mutant analysis shows that alanine racemases from *Pseudomonas aeruginosa* and *Escherichia coli* are dimeric. Journal of Bacteriology. 2002; 184(15):4321–4325.

10. Patrick W, Weisner J, Blackburn J. Site-directed mutagenesis of Tyr354 in *Geobacillus stearothermophilus* alanine racemase identifies a role in controlling substrate specificity and a possible role in the evolution of antibiotic resistance. ChemBioChem. 2002; 3(8):789.

11. Wu D, Hu T, Zhang L, Chen J, Du J, Ding J et al. Residues Asp164 and Glu165 at the substrate entryway function potently in substrate orientation of alanine racemase from *E. coli*: Enzymatic characterization with crystal structure analysis. Protein Science. 2008; 17(6):1066–1076.

12. LeMagueres P, Im H, Ebalunode J, Strych U, Benedik M, Briggs J et al. The 1.9 Å crystal structure of alanine racemase from *Mycobacterium tuberculosis* contains a conserved entryway into the active site. Biochemistry. 2005; 44(5):1471–1481.

13. Shaw J, Petsko G, Ringe D. Determination of the structure of alanine racemase from *Bacillus stearothermophilus* at 1.9Å resolution. Biochemistry. 1997; 36(6):1329–1342.

14. Morollo A, PetskoG, Ringe D. Structure of a Michaelis complex analogue: Propionate binds in the substrate carboxylate site of alanine racemase. Biochemistry. 1999; 38(11):3293–3301.

15. Stamper G, Morollo A, Ringe D. Reaction of alanine racemase with 1-aminoethyl phosphonic acid forms a stable external aldimine. Biochemistry. 1999; 38(20):6714–6714.

16. Kobayashi J, Yukimoto J, Shimizu Y, Ohmori T, Suzuki H, Doi K et al. Characterization of *Lactobacillus salivarius* alanine racemase: Short-chain carboxylate-activation and the role of A131. SpringerPlus. 2015; 4(1):639.

17. LeMagueres P, Im H, Dvorak A, Strych U, Benedik M, Krause K. Crystal structure at 1.45 Å resolution of alanine racemase from a pathogenic bacterium, *Pseudomonas aeruginosa*, contains both internal and external aldimine forms. Biochemistry. 2003; 42(50):14752–14761.

18. Couñago R, Davlieva M, Strych U, Hill R, Krause K. Biochemical and structural characterization of alanine racemase from *Bacillus anthracis (Ames)*. BMC Structural Biology. 2009; 9(1):53.

19. Scaletti E, Luckner S, Krause K. Structural features and kinetic characterization of alanine racemase from *Staphylococcus aureus (Mu50)*. Acta Crystallographica Section D Biological Crystallography. 2011; 68(1):82–92.

20. Priyadarshi A, Lee E, Sung M, Nam K, Lee W, Kim E et al. Structural insights into the alanine racemase from *Enterococcus faecalis*. Biochimica et Biophysica Acta (BBA) — Proteins and Proteomics. 2009; 1794(7):1030–1040.

21. Sun X, He G, Wang X, Xu S, Ju J, Xu X. Crystal structure of a thermostable alanine racemase from *Thermoanaerobacter tengcongensis MB4* reveals the role of Gln360 in substrate selection. PLOS ONE. 2015; 10(7):e0133516.

22. Im H, Sharpe M, Strych U, Davlieva M, Krause K. The crystal structure of alanine racemase from *Streptococcus pneumoniae*, a target for structure-based drug design. BMC Microbiology. 2011; 11(1):116.

23. Noda M, Matoba Y, Kumagai T, Sugiyama M. Structural evidence that alanine racemase from a d-cycloserine-producing microorganism exhibits resistance to its own product. Journal of Biological Chemistry. 2004; 279(44):46153–46161.

24. Sharma V, Wang Y, Liu W. Probing the catalytic charge-relay system in alanine racemase with genetically encoded histidine mimetics. ACS Chemical Biology. 2016; 11(12):3305–3309.

25. Sun S, Toney MD. Evidence for a two-base mechanism involving tyrosine-265 from arginine-219 mutants of alanine racemase. Biochemistry. 1999; 38(13):4058–65.

26. Ciustea M, Mootien S, Rosato A, Perez O, Cirillo P, Yeung K et al. Thiadiazolidinones: A new class of alanine racemase inhibitors with antimicrobial activity against methicillin-resistant *Staphylococcus aureus*. Biochemical Pharmacology. 2012; 83(3):368–377.

27. Lee Y, Mootien S, Shoen C, Destefano M, Cirillo P, Asojo O et al. Inhibition of mycobacterial alanine racemase activity and growth by thiadiazolidinones. Biochemical Pharmacology. 2013; 86(2):222–230.

28. Anthony K, Strych U, Yeung K, Shoen C, Perez O, Krause K et al. New classes of alanine racemase inhibitors identified by high-throughput screening show antimicrobial activity against *Mycobacterium tuberculosis*. PLoS ONE. 2011; 6(5):e20374.

29. Prasad R, Singh A, Srivastava R, Hosmane G, Kushwaha R, Jain A. Frequency of adverse events observed with second-line drugs among patients treated for multidrug-resistant tuberculosis. Indian Journal of Tuberculosis. 2016; 63(2):106–114.

30. Yao X, Skjærven L, Grant B. Rapid characterization of allosteric networks with ensemble normal mode analysis. The Journal of Physical Chemistry B. 2016; 120(33):8276–8288.

31. Miller D, Agard D. Enzyme specificity under dynamic control: A normal mode analysis of α-lytic protease. Journal of Molecular Biology. 1999; 286(1):267–278.

32. Glantz-Gashai Y, Meirson T, Samson A. Normal modes expose active sites in enzymes. PLOS Computational Biology. 2016; 12(12):e1005293.

33. Fuglebakk E, Tiwari S, Reuter N. Comparing the intrinsic dynamics of multiple protein structures using elastic network models. Biochimica et Biophysica Acta (BBA) - General Subjects. 2015; 1850(5):911–922.

34. Kim J, Chang H, Na S. Identification of tail binding effect of kinesin-1 using an elastic network model. Biomechanics and Modeling in Mechanobiology. 2015; 14(5):1107–1117.

35. Kim M, Li W, Shapiro B, Chirikjian G. A Comparison between elastic network interpolation and MD Simulation of 16S ribosomal RNA. Journal of Biomolecular Structure and Dynamics. 2003; 21(3):395–405.

36. Kurkcuoglu Z, Bakan A, Kocaman D, Bahar I, Doruker P. Coupling between catalytic loop motions and enzyme global dynamics. PLoS Computational Biology. 2012; 8(9):e1002705.

37. Hetényi C, van der Spoel D. Blind docking of drug-sized compounds to proteins with up to a thousand residues. FEBS Letters. 2006; 580(5):1447–1450.

38. Hassan N, Alhossary A, Mu Y, Kwoh C. Protein-ligand blind docking using QuickVina-W with inter-process spatio-temporal integration. Scientific Reports. 2017; 7(1):15451.

39. Hetényi C, Spoel D. Toward prediction of functional protein pockets using blind docking and pocket search algorithms. Protein Science. 2011; 20(5):880–893.

40. Levy R, Perahia D, Karplus M. Molecular dynamics of an α-helical polypeptide: Temperature dependence and deviation from harmonic behaviour. Proceedings of the National Academy of Sciences. 1982; 79(4):1346–1350.

41. Swaminathan S, Ichiye T, Van Gunsteren W, Karplus M. Time dependence of atomic fluctuations in proteins: analysis of local and collective motions in bovine pancreatic trypsin inhibitor. Biochemistry. 1982; 21(21):5230–5241.

42. Tama F, Sanejouand Y. Conformational change of proteins arising from normal mode calculations. Protein Engineering, Design and Selection. 2001; 14(1):1–6.

43. Fuglebakk E, Echave J, Reuter N. Measuring and comparing structural fluctuation patterns in large protein datasets. Bioinformatics. 2012; 28(19):2431–2440.

44. Asojo, O., Nelson, S., Mootien, S., Lee, Y., Rezende, W., Hyman, D., Matsumoto, M., Reiling, S., Kelleher, A., Ledizet, M., Koski, R. and Anthony, K. (2014). Structural and biochemical analyses of alanine racemase from the multidrug-resistant *Clostridium difficile* strain 630. Acta Crystallographica Section D Biological Crystallography, 70(7), pp.1922–1933.

45. Huang F, Nau W. A Conformational Flexibility Scale for Amino Acids in Peptides. Angewandte Chemie International Edition. 2003; 42(20):2269–2272.

46. Wenthur C, Gentry P, Mathews T, Lindsley C. Drugs for allosteric sites on receptors. Annual review of Pharmacology and Toxicology. 2014; 54(1):165–184.

47. Berman HM, Westbrook J, Feng Z, Gilliland G, Bhat TN, Weissig H, et al. The Protein Data Bank. Nucleic Acids Res. 2000; 28: 235–242.

48. Kelley L, Mezulis S, Yates C, Wass M, Sternberg M. The Phyre2 web portal for protein modeling, prediction and analysis. Nature Protocols. 2015; 10(6):845–858.

49. Webb B, Sali A. Comparative protein structure modeling using MODELLER. Current Protocols in Protein Science. 2016 Nov 1; 86:2.9.1–2.9.37.

50. The UniProt Consortium. UniProt: the universal protein knowledgebase. Nucleic Acids Research. 2017; 45 (D1):D158–D169.

51. Abraham M, Murtola T, Schulz R, Páll S, Smith J, Hess B et al. GROMACS: High performance molecular simulations through multi-level parallelism from laptops to supercomputers. SoftwareX. 2015; 1–2:19–25.

52. Lindorff-Larsen K, Piana S, Palmo K, Maragakis P, Klepeis J, D ror R et al. Improved side-chain torsion potentials for the Amber ff99SB protein force field. Proteins: Structure, Function, and Bioinformatics. 2010; 78(8):1950–8.

53. Hospital A, Andrio P, Fenollosa C, Cicin-Sain D, Orozco M, Gelpí J. MDWeb and MDMoby: an integrated web-based platform for molecular dynamics simulations. Bioinformatics. 2012; 28(9):1278–1279.

54. de Beer T, Berka K, Thornton J, Laskowski R. PDBsum additions. Nucleic Acids Research. 2013; 42(D1):D292–D296.

55. Laskowski R, MacArthur M, Moss D, Thornton J. PROCHECK: a program to check the stereochemical quality of protein structures. Journal of Applied Crystallography. 1993; 26(2):283–291.

56. Colovos C, Yeates T. Verification of protein structures: Patterns of non-bonded atomic interactions. Protein Science. 1993; 2(9):1511–1519.

57. Ramachandran G, Ramakrishnan C, Sasisekharan V. Stereochemistry of polypeptide chain configurations. Journal of Molecular Biology. 1963; 7(1):95–99.

58. Buchan D, Minneci F, Nugent T, Bryson K, Jones D. Scalable web services for the PSIPRED protein analysis workbench. Nucleic Acids Research. 2013; 41(W1):W349–W357.

59. Altschul S. Gapped BLAST and PSI-BLAST: a new generation of protein database search programs. Nucleic Acids Research. 1997; 25(17):3389–3402.

60. Schaffer A. Improving the accuracy of PSI-BLAST protein database searches with composition-based statistics and other refinements. Nucleic Acids Research. 2001; 29(14):2994–3005.

61. Robert X, Gouet P. Deciphering key features in protein structures with the new ENDscript server. Nucleic Acids Research. 2014; 42(W1):W320–W324.

62. Sievers F, Wilm A, Dineen D, Gibson T, Karplus K, Li W et al. Fast, scalable generation of high-quality protein multiple sequence alignments using Clustal Omega. Molecular Systems Biology. 2014; 7(1):539–539.

63. Lindahl E, Azuara C, Koehl P, Delarue M. NOMAD-Ref: visualization, deformation and refinement of macromolecular structures based on all-atom normal mode analysis. Nucleic Acids Research. 2006; 34(Web Server): W52–W56.

64. Grant B, Rodrigues A, ElSawy K, McCammon J, Caves L. Bio3d: an R package for the comparative analysis of protein structures. Bioinformatics. 2006; 22(21):2695–2696.

65. Skjærven L, Yao X, Scarabelli G, Grant B. Integrating protein structural dynamics and evolutionary analysis with Bio3D. BMC Bioinformatics. 2014; 15(1):399.

66. Ichiye T, Karplus M. Collective motions in proteins: A covariance analysis of atomic fluctuations in molecular dynamics and normal mode simulations. Proteins: Structure, Function, and Genetics. 1991; 11(3):205–217.

67. Hinsen K. Analysis of domain motions by approximate normal mode calculations. Proteins: Structure, Function, and Genetics. 1998; 33(3):417–429.

68. Amadei A, Ceruso M, Di Nola A. On the convergence of the conformational coordinates basis set obtained by the essential dynamics analysis of proteins’ molecular dynamics simulations. Proteins: Structure, Function, and Genetics. 1999; 36(4):419–424.

69. Fuglebakk E, Reuter N, Hinsen K. Evaluation of protein elastic network models based on an analysis of collective motions. Journal of Chemical Theory and Computation. 2013; 9(12):5618–5628.

70. Chemicalize—Instant Cheminformatics Solutions [Internet]. Chemicalize.com. 2018 [cited 21 January 2018]. Available from: https://chemicalize.com

71. ChemAxon—Software Solutions and Services for Chemistry & Biology [Internet]. Chemaxon.com. 2018 [cited 21 January 2018]. Available from: http://www.chemaxon.com

72. Mayo S, Olafson B, Goddard W. DREIDING: a generic force field for molecular simulations. The Journal of Physical Chemistry. 1990; 94(26):8897–8909.

73. Trott O, Olson A. AutoDock Vina: Improving the speed and accuracy of docking with a new scoring function, efficient optimization, and multithreading. Journal of Computational Chemistry. 2009; 31(2):455–61.

74. Grosdidier A, Zoete V, Michielin O. SwissDock, a protein-small molecule docking web service based on EADock DSS. Nucleic Acids Research. 2011; 39(suppl):W270–W277.

75. Schneidman-Duhovny D, Inbar Y, Nussinov R, Wolfson H. PatchDock and SymmDock: servers for rigid and symmetric docking. Nucleic Acids Research. 2005; 33(Web Server):W363–W367.

76. Morris G, Huey R, Lindstrom W, Sanner M, Belew R, Goodsell D et al. AutoDock4 and AutoDockTools4: Automated docking with selective receptor flexibility. Journal of Computational Chemistry. 2009; 30(16):2785–2791.

77. Brooks B, Brooks C, Mackerell A, Nilsson L, Petrella R, Roux B et al. CHARMM: The biomolecular simulation program. Journal of Computational Chemistry. 2009; 30(10):1545–1614.

78. Haberthür U, Caflisch A. FACTS: Fast analytical continuum treatment of solvation. Journal of Computational Chemistry. 2008; 29(5):701–715.

79. Salentin S, Schreiber S, Haupt V, Adasme M, Schroeder M. PL I P: fully automated protein–ligand interaction profiler. Nucleic Acids Research. 2015; 43(W1):W443–W447.

80. Delano W. The PyMOL molecular graphics system. Open source version 1.3. Schrödinger, LLC; 2011.

81. PyMOL Molecular Graphics System — Browse /pymol at SourceForge.net [Internet]. Sourceforge.net. 2018 [cited 21 January 2018]. Available from: https://sourceforge.net/projects/pymol/files/pymol/

82. Dundas J, Ouyang Z, Tseng J, Binkowski A, Turpaz Y, Liang J. CASTp: computed atlas of surface topography of proteins with structural and topographical mapping of functionally annotated residues. Nucleic Acids Research. 2006; 34(Web Server):W116–W118.

83. Laskowski R, Swindells M. LigPlot+: Multiple ligand–protein interaction diagrams for drug discovery. Journal of Chemical Information and Modeling. 2011; 51(10):2778–2786.

